# NK2R signaling governs intestinal lipid mobilization and mucosal inflammation

**DOI:** 10.1101/2025.11.21.689675

**Authors:** Pedro A. Perez, Chung-Chih Liu, Alessandra Ferrari, Nicole Littlejohn, John Paul Kennelly, Emma Robinson, Vân T B Nguyen-Tran, Jon Athanacio, Sean B Joseph, Zaid Amso, Peter Tontonoz, Supriya Srinivasan

**Affiliations:** Department of Neuroscience and Dorris Neuroscience Center, The Scripps Research Institute, 10550 North Torrey Pines Road, La Jolla, CA 92037, USA; Department of Pathology and Laboratory Medicine, University of California, Los Angeles; Los Angeles, CA 90095, USA; Department of Biological Chemistry, University of California, Los Angeles; Los Angeles, CA 90095, USA; Department of Nutritional Sciences and Toxicology, University of California, Berkeley; Berkeley, CA 94720, USA; Howard Hughes Medical Institute, Chevy Chase, MD, USA; Calibr at Scripps Research Institute, 11119 North Torrey Pines Road, La Jolla, CA 92037, USA

## Abstract

Neuropeptidergic control of lipid metabolism is conserved and increasingly implicated in metabolic diseases, but receptor-level mechanisms remain unclear. Here we identify the neurokinin-2 receptor (NK2R) as a central node linking tachykinin signals to intestinal lipid mobilization, epithelial composition, and mucosal inflammation. Across complementary genetic and pharmacological perturbations, modulation of NK2R drives bidirectional effects. Loss or blockade of NK2R increases postprandial triglyceridemia and expands intestinal lipid stores, whereas agonism suppresses chylomicron output, reduces adiposity, and improves glycemia in diet-induced obesity. Transcriptomic and cellular analyses indicate coordinated upregulation of lipid-metabolic programs with a concomitant dampening of immune pathways in the absence of NK2R, accompanied by sex-specific remodeling of secretory lineages and male-biased protection from colitis. NK2R signaling also shaped the fecal microbiota in a genotype- and diet-dependent manner, highlighting crosstalk among neuropeptide signaling, epithelial physiology, and host-microbe interactions. These findings position NK2R as a molecular switch for intestinal lipid handling and mucosal inflammation and suggest that NK2R-targeted agonists or antagonists could be deployed as context- and sex-dependent therapeutic strategies for metabolic disease and inflammatory bowel disease.

## INTRODUCTION

Neuropeptidergic control of lipid metabolism is increasingly recognized as a conserved feature of energy homeostasis^1–7^. Using the *C. elegans* model system, we had previously delineated a brain-to-gut neuroendocrine axis in which the sensory neuron-derived neuropeptide FLP-7 activates the intestinal G protein-coupled receptor NPR-22 to stimulate fat mobilization via induction of the adipocyte triglyceride lipase ATGL-1 and trigger fat loss via mitochondrial β-oxidation and increased energy expenditure^2,8^. Phylogenetic analyses show that FLP-7 sequence resembles the mammalian tachykinin peptide family, and NPR-22 is the *C. elegans* ortholog of the mammalian neurokinin-2 receptor (NK2R)^2^. We have established NPR-22 as both necessary and sufficient for driving fat loss in the worm intestine^2^. Taken together, these findings are consistent with a conserved mammalian tachykinin-NK2R axis controlling lipid mobilization, prompting our examination of mammalian tachykinins and their receptors.

The mammalian tachykinins, including substance P, neurokinin A, and neurokinin B, constitute a conserved neuropeptide family that has historically been known to mediate nociception, inflammation, cancer progression, and gastrointestinal functions^9–13^. These ligands signal through three G protein-coupled receptors (GPCRs): neurokinin-1 receptor (NK1R), neurokinin-2 receptor (NK2R), and neurokinin-3 receptor (NK3R), which are preferentially activated by substance P, neurokinin A, and neurokinin B, respectively. Prior studies have established NK1R as a key regulator of pain, inflammation, and immune responses^14–17^. NK1R antagonists have demonstrated therapeutic potential by attenuating blood-brain barrier dysfunction, cerebral edema, and pro-inflammatory cytokine levels in traumatic brain injury models^18–21^. In contrast, NK3R has been extensively studied in reproductive biology and neurological disorders^11,22,23^. The neurokinin B-NK3R signaling is critical for hypothalamic-pituitary-gonadal axis function, with an NK3R antagonist now clinically deployed to relieve menopausal vasomotor symptoms^24,25^. Beyond reproduction, NK3R has also been implicated in addiction, pain, and psychiatric disorders^22,26–28^.

By comparison, NK2R regulates diverse physiological processes, including smooth muscle contraction, nociception, intestinal fluid secretion, and reproductive functions^29–34^. Although NK2R is expressed in multiple tissues, including the gastrointestinal tract, lung, and immune system^35–41^, its specific roles in metabolism and immunity remain poorly defined. A recent study^42^ reported that the administration of a selective NK2R peptide agonist increases energy expenditure, reduces fat mass, and suppresses appetite, highlighting its therapeutic potential for cardiometabolic diseases, including obesity and diabetes. However, the molecular mechanisms engaged by NK2R agonism are largely unknown.

## RESULTS

### Expression pattern of Tachykinin Receptor 2 (*Tacr2*)

Our prior work in *C. elegans* showed that the CeNK2R gene *npr-22* is expressed in a few pairs of head neurons and in the intestine, but not in the muscle or hypodermal tissues^2^ (Fig. 1a). We examined *Tacr2* expression in mammalian species: the Human Protein Atlas^43,44^ RNA-seq datasets revealed that human *TACR2* (*hTACR2*) is preferentially co-expressed with genes enriched in the intestinal “digestion” cluster (Fig. 1b). In contrast, *hTACR1* and *hTACR3* clustered with transcripts characteristic of connective tissue and retina, respectively (Extended Data Fig. 1a). Consistent with these patterns, quantitative PCR across mouse small-intestinal segments showed that *Tacr2* mRNA levels exceeded those of *Tacr1* and *Tacr3* along the entire proximal-distal axis (Fig. 1c). Immunofluorescence localized the NK2R protein to the basolateral membrane of mouse intestinal epithelial cells (Fig. 1d), indicating a potential role in responsiveness to internal physiological signals rather than luminal nutrients. These results are consistent with our previous findings in *C. elegans*^2^, in which NPR-22, the invertebrate ortholog of NK2R, is similarly expressed in intestinal cells and responds to internal signals from sensory neurons.

**Figure 1.**
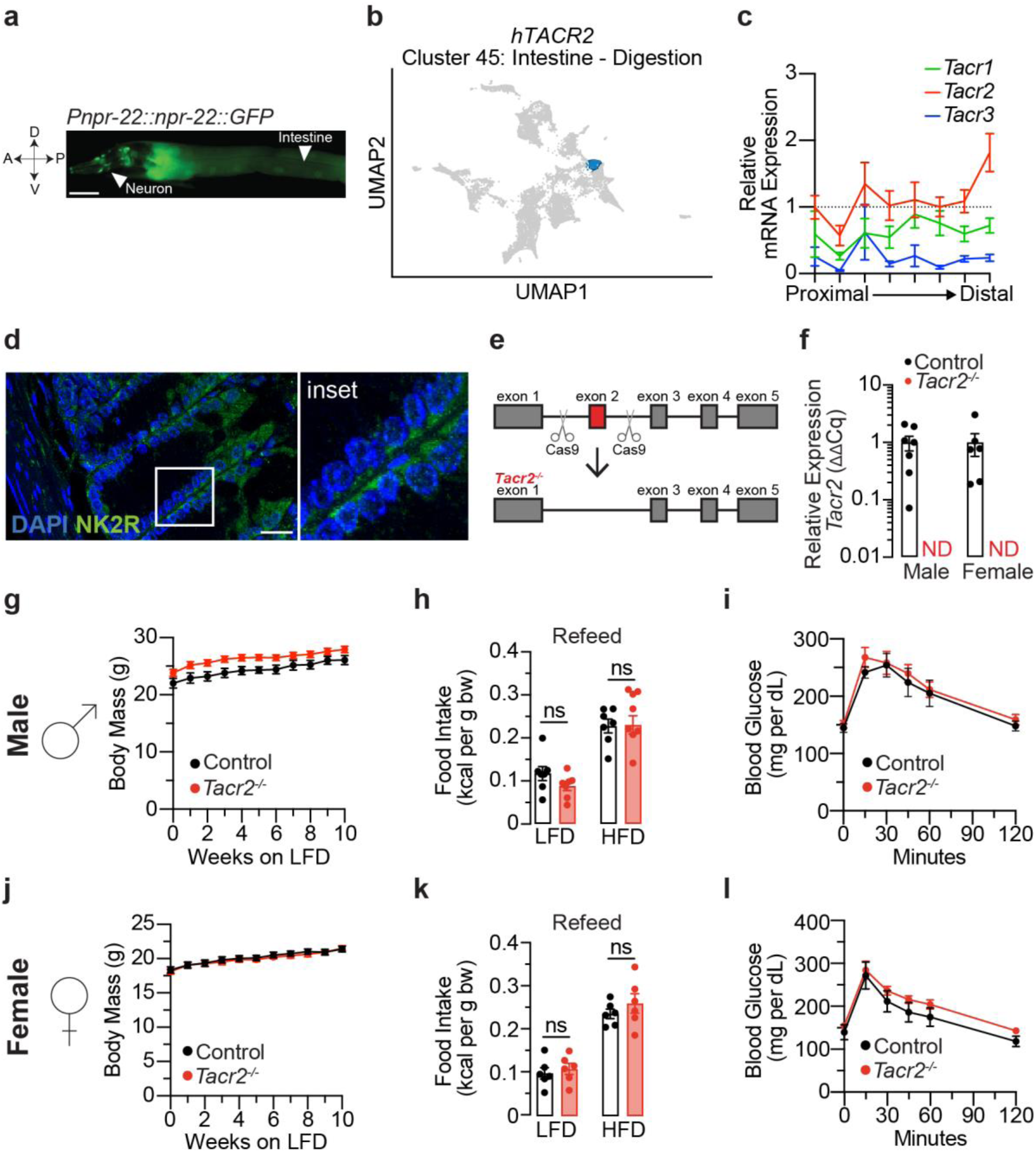
Tachykinin receptor 2 tissue expression. **a**, Fluorescent image of a transgenic *C. elegans* expressing *npr-22::GFP* under the control of the endogenous *npr-22* promoter. GFP expression was detected in the intestinal cells and in several pairs of neurons in the head. A, anterior; P, posterior; V, ventral; D, dorsal. Scale bar, 50 μm. **b**, UMAP visualization depicting gene clusters based on mRNA expression profiles from The Human Protein Atlas^44^ (HPA). The cluster containing *hTacr2* (cluster 45, comprising 300 genes) is highlighted in blue. Functional annotation provided by HPA characterizes the shared specificity (Intestine) and biological function (Digestion) of genes within this cluster. **c**, The gene expression levels of *Tacr1*, *Tacr2*, and *Tacr3* were assessed throughout the entire mouse small intestine using qPCR. The intestine was divided into eight equal segments, each measuring approximately 4-5 cm. *Actb* was used as the housekeeping gene. Data are presented as the fold change relative to the expression level of *Tacr2* in the first segment ± SEM. n = 6 biological replicates. **d**, Fluorescent image of mouse jejunum stained with NK2R antibody (green) and DAPI (blue). Scale bar: 20 μm. Inset: magnified view of the region enclosed by the white line in the fluorescent image. The jejunum was harvested from male C57BL6/J mice (8 weeks old, low-fat diet). **e**, The strategy of CRISPR-Cas9 mediated genome editing, depicting the genomic region of *Tacr2* and the locations of the Cas9 cutting sites for *Tacr2^-/-^*. The deleted exon 2 is marked in red. **f**, qPCR analysis of *Tacr2* mRNA expression in intestinal epithelial RNA isolated from male and female control and *Tacr2^-/-^* mice. n = 7 male control, n = 6 female control, n = 6 male *Tacr2^-/-^*, n= 6 female *Tacr2^-/-^*. **g**,**j**, Body weights of male (**g**) and female (**j**) *Tacr2^-/-^* and control mice (8-12 weeks old) were monitored weekly over a 10-week period while maintained on a low-fat diet (LFD). n = 8 control and n = 12 *Tacr2^-/-^*mice (**g**); n = 11 control and n = 13 *Tacr2^-/-^* mice (**j**). **h**,**k**, Following an overnight fast, male (**h**) and female (**k**) mice were re-fed with either LFD or HFD, and cumulative food intake was measured over a 4 hr refeeding period. n = 7 control (LFD), n = 7 *Tacr2^-/-^* (LFD), n = 7 control (HFD), and n = 8 *Tacr2^-/-^* (HFD) mice **(h)**; n = 6 per group (**k**). **i,l,** Glucose tolerance tests were performed in week 10 in male **(i)** and female (**l**) mice. n = 5 control and n = 6 *Tacr2^-/-^*mice per group. Data was analyzed using an unpaired Student’s t-test (two-tailed) (**f,h,k**) or two-way ANOVA with repeated measures (**g,i,j,l**). Data are shown as mean ± SEM. ^ns^p>0.05. Black bars/lines represent control mice; red bars/lines represent *Tacr2^-/-^* mice.

### Generation of *Tacr2*-null mice and analysis of gene expression in response to dietary challenges

To investigate NK2R function *in vivo*, we generated global *Tacr2*-null (*Tacr2-/-*) mice by CRISPR-Cas9 gene editing (Fig. 1e). qPCR of jejunal epithelium confirmed loss of *Tacr2* transcripts (Fig. 1f) without compensatory up-regulation of *Tacr1* or *Tacr3* (Extended Data Fig. 1b). *Tacr2-/-* mice of both sexes were grossly normal, displaying wild-type body mass, re-feeding behavior and glucose tolerance (Fig. 1g-l).

Given the role of the *C. elegans* ortholog NPR-22 in intestinal lipid metabolism^2^, we wished to determine whether NK2R modulates mammalian intestinal lipid handling. *Tacr2-/-* and wild-type littermates were maintained for one week on either a low-fat diet (LFD) or a western-style high-fat diet (HFD) before jejunal epithelial RNA was profiled by RNA-seq (Fig. 2a). Body mass remained comparable between *Tacr2*-/- and wild-type littermates of both sexes on either LFD or HFD (Extended Data Fig. 2a, b).

**Figure 2.**
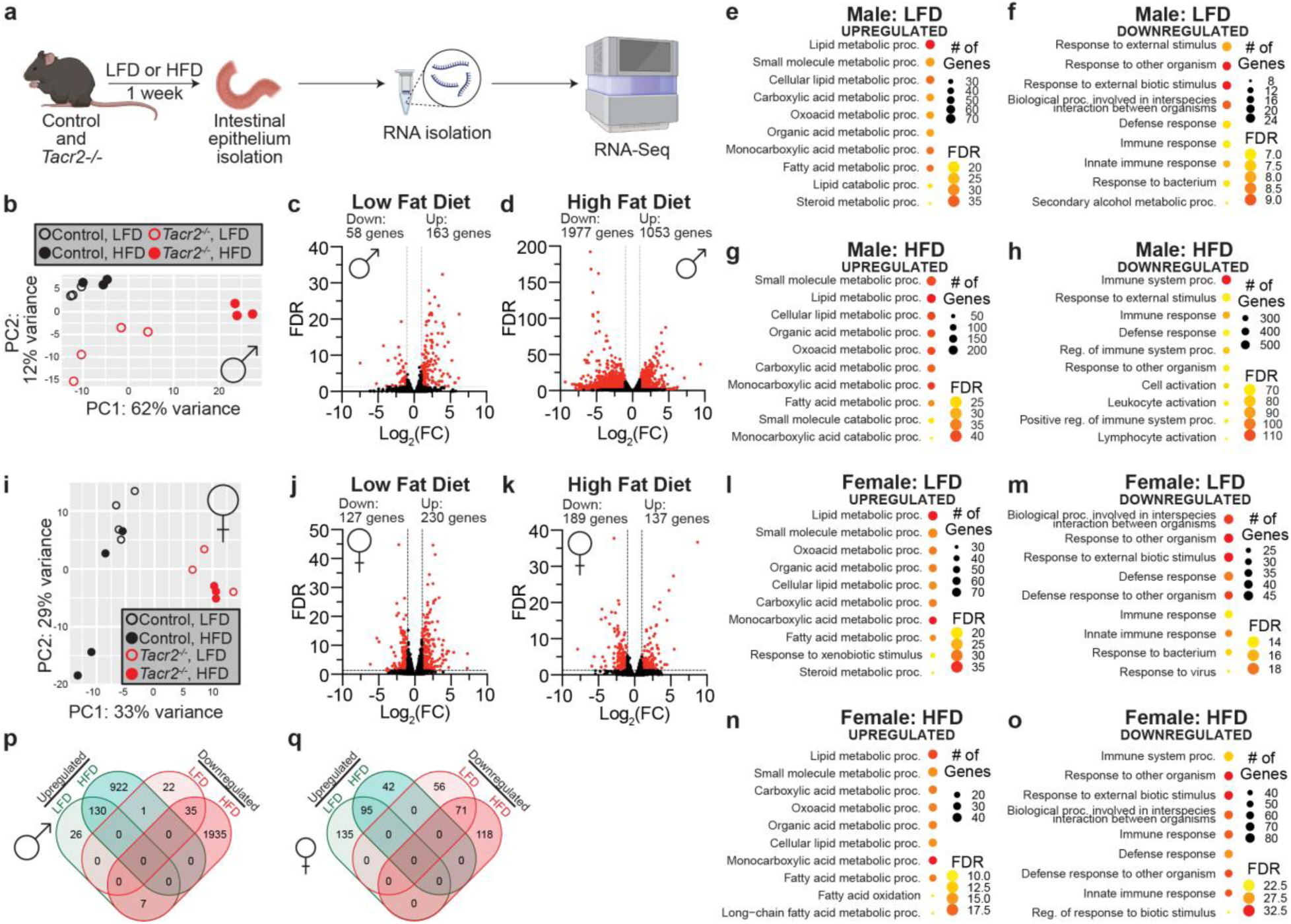
Sex-specific transcriptional responses to high-fat diet in *Tacr2^-/-^* mice. **a**, Schematic of the experimental workflow. Control and *Tacr2^-/-^* mice were fed a low-fat diet (LFD) or high-fat diet (HFD) for 1 week before sacrifice. Following tissue collection, the upper intestinal epithelial layer was isolated, and total RNA was extracted for subsequent bulk RNA sequencing. **b**, Principal component analysis (PCA) of the RNA-Seq data from male control and *Tacr2^-/-^*mice on LFD or HFD. n = 3 control (LFD), n = 3 control (HFD), n = 4 *Tacr2^-/-^*(LFD), n = 3 *Tacr2^-/-^* (HFD). **c**,**d**, Volcano plots of differentially expressed genes in male *Tacr2^-/-^* versus control mice under LFD (**c**) or HFD (**d**) conditions. **e**-**h**, Gene ontology (GO) enrichment analyses of genes differentially expressed in male *Tacr2^-/-^* mice versus controls on LFD (**e**, upregulated; **f**, downregulated) or HFD (**g**, upregulated; **h**, downregulated). Bubble size indicates the number of genes associated with each GO term, and color represents the false discovery rate (FDR). **i**, Principal component analysis of RNA-Seq data from female control and *Tacr2^-/-^* mice on LFD or HFD. n = 4 control (LFD), n = 4 control (HFD), n = 3 *Tacr2^-/-^*(LFD), n = 3 *Tacr2^-/-^* (HFD). **j**,**k**, Volcano plots of differentially expressed genes in female *Tacr2^-/-^* versus control mice under LFD (**j**) or HFD (**k**) conditions. **l**-**o**, Gene ontology enrichment analyses of genes differentially expressed in female *Tacr2^-/-^*mice versus controls on LFD (**l**, upregulated; **m**, downregulated) or HFD (**n**, upregulated; **o**, downregulated). Bubble size and color reflect gene count and FDR, respectively. **p**,**q**, Venn diagrams showing overlap of differentially expressed genes (adjusted p-value < 0.05 and |log_2_ fold change| > 1.0) between LFD and HFD conditions in male (**p**) and female (**q**) datasets. Numbers indicate the number of shared or unique genes between conditions.

In males, principal component analysis (PCA) revealed distinct clustering patterns: the wild-type mice clustered closely irrespective of diet; *Tacr2*-/- mice fed a LFD formed a separate cluster from wild-types; *Tacr2*-/- mice fed a HFD exhibited markedly different clustering compared to all other groups (Fig. 2b). Only 221 genes were differentially expressed (DE) between genotypes on LFD, whereas 3030 genes were DE on HFD (Fig. 2c, d), suggesting that loss of *Tacr2* selectively sensitizes the male intestinal epithelium to dietary fat. Interestingly, Gene Ontology (GO) enrichment linked up-regulated transcripts to lipid-metabolic pathways and down-regulated transcripts to immune and inflammatory processes (Fig. 2e-h), concordant with prior *C. elegans* work that demonstrates a role for NK2R in intestinal lipid metabolism^2,8^.

In females, the PCA was driven principally by genotype rather than diet (Fig. 2i). The number of DE genes remained similar under LFD and HFD (Fig. 2j, k), yet functional annotations mirrored those in males: lipid metabolism among up-regulated genes and immune response among down-regulated genes (Fig. 2l-o). Overlap between diet-specific DE gene sets was modest in males but substantial in females (Fig. 2p, q), indicating a differential response in males to dietary fat in the absence of NK2R.

### NK2R governs epithelial lineage allocation and intestinal inflammation

Interestingly, genes downregulated in *Tacr2-/-* jejunum of both sexes were enriched for GO terms linked to immune response, immune regulation and leukocyte activation (Fig. 2f, h, m, o), indicating an overall dampening of mucosal immunity. Bisque deconvolution^45^ of bulk RNA-seq revealed a basal shift in epithelial lineages in male *Tacr2-/-* mice (Fig. 3a). Tuft and transit-amplifying (TA) cell signatures were elevated, whereas stem-cell and goblet-cell signatures were reduced; HFD further exacerbated these baseline genotype differences, notably by reducing the relative proportion of enterocytes. Paneth cell signatures declined after HFD in both genotypes, consistent with previous studies^46,47^, while endocrine cell abundance was unchanged. In contrast, female mice showed only minor alterations (Extended Data Fig. 3a); in *Tacr2-/-* females, HFD lowered stem-cell and enterocyte signatures and increased TA cells.

**Figure 3.**
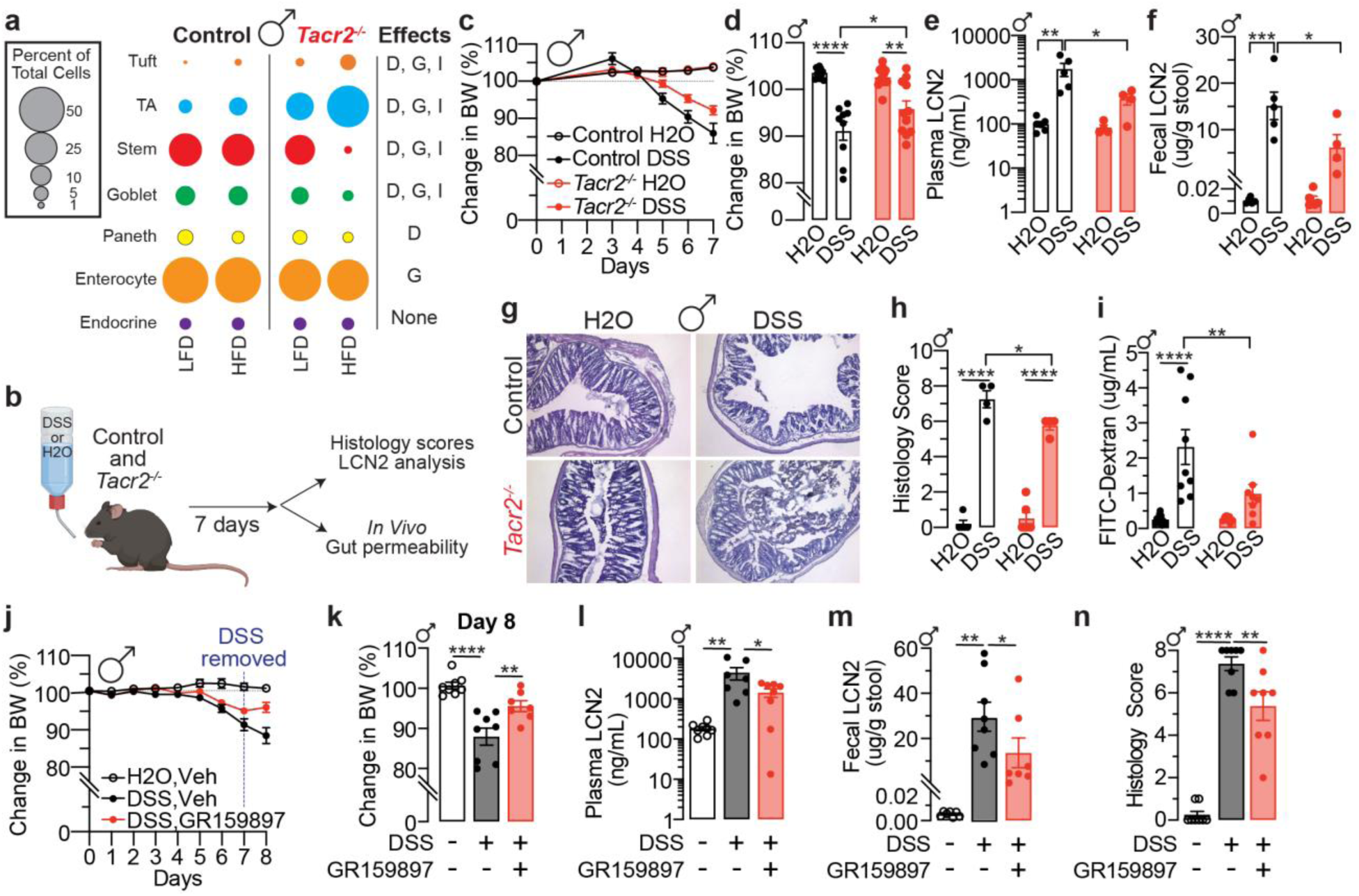
NK2R inhibition attenuates DSS-induced weight loss and intestinal inflammation in male mice. **a**, Bulk RNA-Seq data were integrated with publicly available single-cell RNA-Seq datasets to estimate cell-type composition in the small intestine of male mice. Differences in estimated cell-type proportions were analyzed based on diet (D), genotype (G), or their interaction (I). **b**, Schematic representation of the experimental workflow for the DSS-induced colitis model. Male and female mice received DSS solution (3% for males, 5% for females) or vehicle (drinking water) ad libitum for 7 consecutive days. Following treatment, tissues were collected for downstream analyses or subjected to an *in vivo* gut permeability assay. **c**, Daily monitoring of body weights in male control and *Tacr2^-/-^* mice treated with DSS or vehicle over a 7-day period. n = 5 control (H_2_O), n = 5 control (DSS), n = 6 *Tacr2^-/-^*(H_2_O), and n = 6 *Tacr2^-/-^* (DSS). **d**, Comparison of body weight changes from baseline (day 0) to day 7, derived from the data shown in panels (**c**)**. e**,**f**, Lipocalin-2 (LCN2) concentrations in blood (**e**) and feces (**f**) from male control and *Tacr2^-/-^*mice after 7 days of DSS or vehicle treatment, as determined by ELISA. n = 6 control (H_2_O), n = 5 control (DSS), n = 4 *Tacr2^-/-^* (H_2_O), and n = 4 *Tacr2^-/-^* (DSS) for (**e**); n = 5 control (H_2_O), n = 5 control (DSS), n = 5 *Tacr2^-/-^* (H_2_O), and n = 4 *Tacr2^-/-^* (DSS) for (**f**). **g**, Colon tissues from male control and *Tacr2^-/-^*mice treated with DSS or vehicle for 7 days were collected, sectioned, and stained with hematoxylin and eosin (H&E). Representative images of stained colon sections are shown. **h**, Representative histological images (**g**) were evaluated and scored according to the degree of tissue damage. n = 5 control (H_2_O), n = 4 control (DSS), n = 6 *Tacr2^-/-^* (H_2_O), and n = 4 *Tacr2^-/-^*(DSS). **i**, Gut permeability in male control and *Tacr2^-/-^*mice following 7 days of DSS or vehicle treatment was assessed by oral gavage of 4 kDa FITC-dextran. Blood was collected 4 hours post-gavage, and FITC fluorescence was measured in plasma to quantify gut barrier integrity. n = 10 control (H_2_O), n = 9 control (DSS), n = 9 *Tacr2^-/-^* (H_2_O), and n = 8 *Tacr2^-/-^* (DSS). **j**, Daily monitoring of body weights in male wild-type mice treated with DSS or vehicle for 7 days, in combination with daily injections of the NK2R antagonist GR159897 (2.5 mg/kg) or vehicle (PBS). DSS administration was discontinued on day 7. n = 8 (H_2_O+Vehicle), n = 8 (DSS+Vehicle), n = 7 (DSS+GR159897). **k**, Comparison of body weight changes from baseline (day 0) to day 8, derived from the data shown in panel (**j**). **l**,**m**, Lipocalin-2 (LCN2) concentrations in blood (**l**) and feces (**m**) from male wild-type mice with indicated treatment on day 8, as determined by ELISA. n = 8 (H_2_O+Vehicle), n = 7 (DSS+Vehicle), and n = 8 (DSS+GR159897) for (**l**); n = 7 (H_2_O+Vehicle), n = 8 (DSS+Vehicle), and n = 8 (DSS+GR159897) for (**m**). **n**, Colon tissues from male wild-type mice with indicated treatment on day 8 were collected, sectioned, and stained with hematoxylin and eosin (H&E). Representative histological images were evaluated and scored according to the degree of tissue damage. n = 8 (H_2_O+Vehicle), n = 8 (DSS+Vehicle), and n = 8 (DSS+GR159897). Statistical analysis was performed using one-way or two-way ANOVA followed by Holm-Sidak post hoc tests. Data are shown as mean ± SEM, ****p < 0.0001, ***p < 0.001, **p < 0.01, *p < 0.05. Black bars/lines represent control mice; red bars/lines represent *Tacr2^-/-^* or GR159897-treated mice.

Building on the transcriptional changes in immune-related genes observed in *Tacr2-/-* mice, we next assessed functional consequences of *Tacr2* loss *in vivo* by inducing colitis in mice using the well-established dextran sodium sulfate (DSS) model^48^ (Fig. 3b). Male *Tacr2-/-* mice were protected from the deleterious effects of induced colitis, exhibiting attenuated weight loss, reduced plasma and fecal lipocalin-2 (LCN2) concentration (a marker of intestinal inflammation^49^), ameliorated histopathology with preserved architecture and less immune-cell infiltration. In a second experiment performed under identical conditions, we observed improved epithelial barrier function indicated by reduced FITC-dextran permeability via the intestinal lumen (Fig. 3c-i). In contrast, female *Tacr2-/-* mice showed no protection in weight loss or plasma LCN2, increased fecal LCN2, and histological injury comparable to controls, although barrier integrity was partially maintained (Extended Data Fig. 3b-h). Of note, male *Tacr2*-/- mice exhibited increased numbers of differentially expressed genes following HFD, an inflammatory insult^50–53^, and demonstrated colitis protection, whereas female *Tacr2*^-/-^mice lacked this expanded gene expression response and did not exhibit protection against DSS-induced colitis. Together, these data indicate a sex-dependent transcriptional and functional responses to inflammatory challenge in the absence of NK2R.

Prompted by this result, we tested whether acute receptor antagonism via a selective NK2R antagonist would phenocopy the genetic absence, and help ascertain translational tractability. Single-dose pharmacological inhibition corroborated the genetic findings: daily administration of the selective NK2R antagonist GR159897 during DSS exposure mitigated weight loss, suppressed plasma and fecal LCN2, and improved histological scores in males (Fig. 3j-n). Together, these data identify NK2R as a sex-dependent regulator of epithelial lineage allocation and susceptibility to mucosal injury.

### Role of NK2R in Lipid Metabolism

Guided by our jejunal epithelial RNA-seq results, we next wished to determine the role of NK2R in lipid metabolism. Genes up-regulated in the *Tacr2-/-* jejunal epithelium were enriched for GO categories related to lipid and small-molecule metabolism, and this signature persisted and was further amplified under HFD challenge, indicating enhanced sensitivity to luminal lipid load. Because enterocyte lipid uptake culminates in cytosolic lipid droplet (LD) storage before export as chylomicrons, we profiled a curated set of 116 LD-associated genes^54^ to examine whether NK2R modulates LD dynamics. In *Tacr2-/-* males, hierarchical clustering segregated HFD-fed *Tacr2-/-* samples from all other groups, whereas wild-type (WT) controls clustered together irrespective of diet (Fig. 4a). *Tacr2-/-* females also exhibited broad up-regulation of LD genes, although clustering by genotype and diet was less pronounced (Extended Data Fig. 5a).

**Figure 4.**
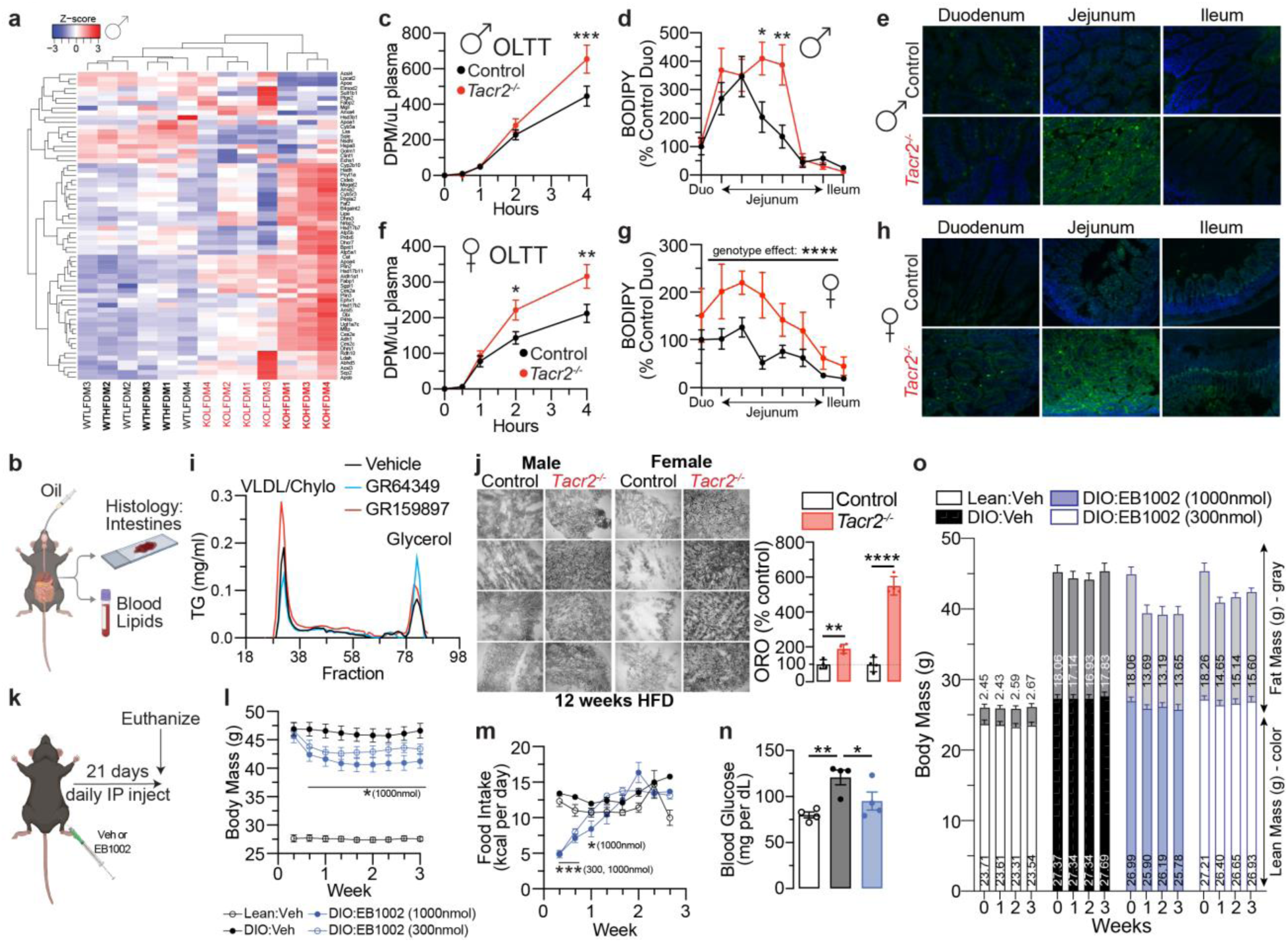
*Tacr2^-/-^*mice exhibit enhanced intestinal lipid absorption. **a**, Heat maps display Z-scores of normalized expression levels for genes encoding lipid droplet-associated proteins in male mice across experimental conditions. Hierarchical clustering of individual samples (top dendrogram) and genes (left dendrogram) is shown, with branch lengths reflecting the similarity between clusters. Wild-type control (WT) samples are labeled in black text, *Tacr2^-/-^*(KO) samples in red text, and samples from mice on a high-fat diet (HFD) are indicated in bold. **b**, Schematic representation of the experimental workflow for assessing intestinal lipid absorption. Mice were administered an oral gavage of oil containing either 14C-triolein (**c**,**f**) or C16-BODIPY (**d**,**e**,**g**,**h**), followed by quantification of lipids in intestinal tissue and blood. **c**,**f**, Oral lipid tolerance test (OLTT) was performed in male (**c**) and female (**f**) control and *Tacr2^-/-^* mice. Blood lipid levels were measured over a 4-hour period following oral gavage and expressed as disintegrations per minute (DPM) per μL of plasma. n = 8 control males, n = 7 *Tacr2^-/-^*males for (**c**); n = 7 control females, n = 9 *Tacr2^-/-^* females for (**f**). **d**,**g**, BODIPY fluorescence was measured in intestinal sections from male (**d**) and female (**g**) control and *Tacr2^-/-^*mice following oral gavage with oil containing C16-BODIPY. n = 5 control males, n = 5 *Tacr2^-/-^* males (**d**); n = 6 control females, n = 7 *Tacr2^-/-^* females (**g**). **e**,**h**, Representative fluorescence images of intestinal sections from male (**e**) and female (**h**) mice corresponding to the samples shown in panels (**d**) and (**g**), respectively. **i**, Triglyceride levels were measured in each plasma fraction from wildtype mice fed a HFD for 10 weeks and pretreated with LPL inhibitor and either vehicle, the NK2R agonist GR64349, or NK2R antagonist GR159897 prior to oral oil gavage. n=8 mice per treatment group. **j**, Representative images and quantification of Oil Red O (ORO)-stained intestinal sections from male and female control and *Tacr2^-/-^* mice following 12 weeks of HFD feeding. n = 4 control males, n = 4 *Tacr2^-/-^* males, n = 4 control females, n = 4 *Tacr2^-/-^*females. **k**, Schematic representation of the experimental workflow for assessing the effects of NK2R agonism on body mass. Diet-induced obese (DIO) wild-type mice were administered daily intraperitoneal injections of EB1002 or vehicle for 21 consecutive days. **l**,**m**, Body weights (**l**) and food intake (**m**) were monitored over a 3-week period in lean and diet-induced obese (DIO) mice treated with vehicle, high-dose EB1002 (1000 nmol), or low-dose EB1002 (300 nmol). n = 8 (lean: vehicle), n = 7 (DIO: vehicle), n = 8 (DIO: EB1002, 1000nmol) and n = 8 (DIO: EB1002, 300nmol). **n**, The fasted blood glucose levels were determined in lean and diet-induced obese (DIO) mice treated with vehicle or high-dose EB1002 (1000 nmol). n = 4 (lean: vehicle), n = 4 (DIO: vehicle), and n = 4 (DIO: EB1002, 1000nmol). **o**, Lean mass and fat mass were measured over 3 weeks in lean and diet-induced obese (DIO) mice treated with vehicle, high-dose EB1002 (1000 nmol), or low-dose EB1002 (300 nmol). n = 8 (lean: vehicle), n = 7 (DIO: vehicle), n = 8 (DIO: EB1002, 1000nmol) and n = 8 (DIO: EB1002, 300nmol). Statistical analyses were performed using two-way ANOVA with repeated measures (**c**,**d**,**f**,**g**,**l**,**m**), unpaired two-tailed Student’s t-test (**b**,**j**), one-way ANOVA with Holm-Sidak post hoc analysis (**n**), or two-way ANOVA with Holm-Sidak post hoc analysis (**o**). Data are presented as mean ± SEM. ****p < 0.0001, ***p < 0.001, **p < 0.01, *p < 0.05, ^ns^p>0.05. Black bars/lines indicate control mice, red bars/lines indicate *Tacr2^-/-^* mice and blue bars/lines indicate NK2R agonist-treated mice.

The enrichment of lipid-metabolic pathways and coordinated upregulation of the enterocyte LD program in the *Tacr2-/-* jejunal epithelium suggested that NK2R constrains the absorptive phase of dietary lipid handling and therefore regulates luminal lipid uptake. Following oil gavage, *Tacr2-/-* mice exhibited normal gastrointestinal transit (Extended Data Fig. 4a) but displayed a higher post-prandial triglyceride excursion (oral lipid-tolerance test, OLTT) and increased jejunal lipid retention 4 h after dosing in both sexes (Fig. 4b-h). Importantly, the OLTT reports the appearance of intestinally derived triglycerides in plasma. Thus, the enhanced excursion in *Tacr2-/-* mice indicates increased intestinal lipid entry into the circulation. The concurrent increase in jejunal lipid stores suggests that luminal uptake and intracellular storage of lipids are augmented, consistent with the observed induction of the LD program. Furthermore, pharmacological modulation recapitulated these effects: in diet-induced obese mice pre-treated with tyloxapol to inhibit lipoprotein lipase (LPL) and to block peripheral triglyceride clearance, the NK2R antagonist GR159897 elevated chylomicron/very-low-density lipoprotein (VLDL) triglycerides, whereas the NK2R agonist GR64349 produced the opposite outcome (Fig. 4i). Chronic HFD feeding for 12 weeks markedly expanded intestinal epithelial LD stores in *Tacr2-/-* mice of both sexes relative to wild-type controls (∼2-fold in males, ∼5-fold in females; Fig. 4j) without altering food intake, intestinal motility, metabolic rates, or locomotor activity (Extended Data Figs. 4b-k, 5b-k). Together, these data support a model in which NK2R signaling tonically regulates lipid metabolism in enterocytes by tuning luminal lipid uptake, intracellular LD storage and chylomicron secretion.

Since adipose tissue is one of the major depots for chylomicron-derived fatty acids^55^, the intestinal phenotypes above predict altered lipid delivery to fat depots. Moreover, *Tacr2* expression in white adipose tissue (WAT) (Extended Data Fig. 6a) raises the possibility of adipose-autonomous NK2R actions independent of the intestine. We therefore asked whether loss of NK2R remodels adipose transcriptional programs and wished to distinguish secondary consequences of increased intestinal export from adipose-intrinsic NK2R effects. *Tacr2* is expressed in WAT, with comparable expression in both adipocyte and mesenchymal cell fractions (Extended Data Fig. 6a). To assess potential transcriptional changes, we performed RNA sequencing with perigonadal WAT at early (1 week) and chronic (12 weeks) time points after HFD exposure. (Extended Data Fig. 6b). Remarkably, the PCA revealed no genotype-dependent clustering in either male or female mice at either time point (Extended Data Fig. 6c-j). Furthermore, the number of differentially expressed genes between *Tacr2-/-* and wild-type mice was negligible, indicating that *Tacr2* deletion does not appreciably alter the WAT transcriptome under HFD conditions. Together, these results suggest that, despite robust intestine-specific effects on lipid metabolism, the absence of NK2R has a limited impact on adipose transcriptional remodeling during diet-induced obesity.

Given that acute pharmacological activation of NK2R lowered chylomicron-associated triglycerides, we next asked whether sustained NK2R agonism alters whole-body energy balance. Diet-induced obese (DIO) mice received daily intraperitoneal injections of the NK2R-selective peptide agonist EB1002 (1000 nmol per kg) for 21 days (Fig. 4k). EB1002 is a long-acting NK2R agonist recently reported by the Gerhart-Hines group to improve cardiometabolic parameters in DIO models^42^. In our experiments, EB1002 treatment reduced body weight, perigonadal fat mass, and fasting glycemia of DIO mice relative to vehicle controls, concordant with the recent report^42^ (Fig. 4l, n, o). Food intake was transiently suppressed during the first week of treatment but normalized thereafter (Fig. 4m), indicating that the weight loss was not solely attributable to persistent hypophagia. Together with the acute lipid-handling phenotypes, these data position NK2R as a key direct regulator of intestinal lipid mobilization and long-term systemic energy balance.

### Male *Tacr2-/-* mice have altered microbial populations

To determine whether *Tacr2-/-* males are uniquely prone to HFD-induced dysbiosis, fecal bacterial communities were profiled by 16S rRNA gene sequencing. α-diversity, assessed by the Shannon index, was reduced in *Tacr2-/-* mice irrespective of diet (Fig. 5a). β-diversity analysis (weighted UniFrac distances) revealed significant main effects of genotype (PERMANOVA, p=0.002) and diet (p=0.017), as well as a genotype-diet interaction (p=0.001) (Fig. 5b), indicating that both factors independently and synergistically shape the fecal microbiota, with *Tacr2* deletion predisposing to larger diet-driven compositional shifts.

**Figure 5.**
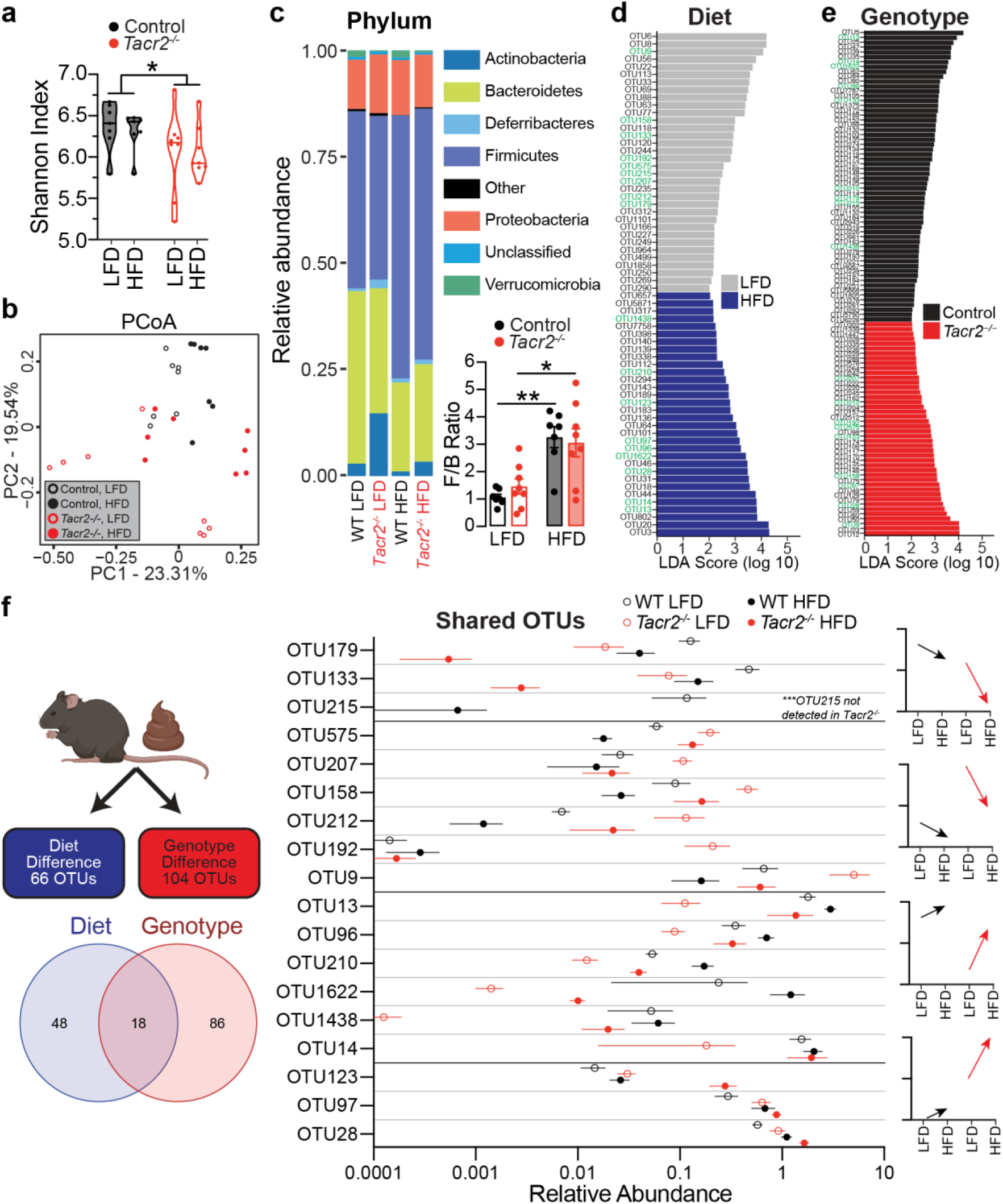
16S rRNA sequencing analysis of gut microbial populations. **a**, Shannon index (α-diversity) of fecal microbial populations in control and *Tacr2^-/-^* mice fed a LFD or HFD. n = 7 control (LFD), n = 7 control (HFD), n = 8 *Tacr2^-/-^* (LFD), n = 8 *Tacr2^-/-^*(HFD). Data were analyzed using two-way ANOVA followed by Holm-Sidak’s post hoc analysis; genotype effect: p = 0.039, diet effect: ns, genotype x diet interaction: ns. **b**, Principal Coordinate Analysis (PCoA) plot based on weighted UniFrac distances illustrating β-diversity of microbial communities in *Tacr2^-/-^* and control mice fed a LFD or HFD. Each point represents a microbial community from an individual sample, color-coded by genotype (*Tacr2^-/-^*, red; control, black) and diet (LFD, open circles; HFD, closed circles). **c**, Microbial populations were sorted at the phylum level across experimental conditions. *Firmicutes*/*Bacteroidet*es (F/B) ratio was calculated in control and *Tacr2^-/-^* mice fed a LFD or HFD. n = 7 control (LFD), n = 8 *Tacr2^-/-^* (LFD), n = 7 control (HFD), n = 8 *Tacr2^-/-^*(HFD). Data were analyzed using two-way ANOVA followed by Holm-Sidak post hoc analysis. Data are shown as mean ± SEM; **p<0.01, *p<0.05. **d**,**e**, Operational Taxonomic Units (OTUs) were analyzed using Linear Discriminant Effect Size (LEfSe) to identify taxa differentially enriched by diet or genotype. **d**, Bar graph showing Linear Discriminant Analysis (LDA) scores for 66 diet-dependent OTUs (34 associated with LFD, 32 associated with HFD). **e**, Bar graph showing LDA scores for 104 genotype-dependent OTUs (60 associated with control mice, 44 associated in *Tacr2^-/-^* mice). **f**, Relative abundances of 18 OTUs shared between the diet- and genotype-dependent analyses. OTUs were filtered using a Linear Discriminant Analysis (LDA) score threshold > 2.0. In panels (**d**) and (**e**), OTUs shared between diet and genotype analyses are indicated in green text.

At the phylum level, HFD feeding increased the relative abundance of *Firmicutes* and concomitantly decreased *Bacteroidetes*, yielding the characteristic rise in the *Firmicutes*:*Bacteroidetes* ratio^46,56,57^ (Fig. 5c). A similar shift was observed in *Tacr2-/-*mice under HFD conditions. Linear discriminant analysis effect size (LEfSe) identified 66 operational taxonomic units (OTUs) that were diet-responsive and 104 that were genotype-responsive (Fig. 5d, e); 18 OTUs were influenced by both variables (Fig. 5f). Notably, OTU 215 was completely absent in *Tacr2-/-* mice, revealing a genotype-specific microbial deficit.

## DISCUSSION

In this report we identify neurokinin-2 receptor (NK2R) signaling as a nexus between intestinal lipid handling, mucosal immunity and host-microbe interactions. *Tacr2* deletion or acute pharmacological antagonism augmented chylomicron export and massively expanded lipid-droplet (LD) stores in jejunal enterocytes, whereas NK2R agonism suppressed post-prandial triglyceridemia and reduced adiposity. These bidirectional effects suggest that NK2R serves as a molecular switch for intestinal lipid mobilization. The downstream cascade by which NK2R transduces neuropeptidergic input to LD dynamics remains to be elucidated; however, transcriptomic profiling of *Tacr2-/-* jejunal epithelium from the present study uncovered the coordinated up-regulation of a core LD gene module, suggesting that tonic NK2R activity normally restrains this program.

Bisque deconvolution revealed that *Tacr2*-null males, but not females, remodeled goblet and tuft cell lineages when challenged with a HFD. These secretory cell types shape barrier function and epithelial sensing of luminal antigens^58,59^ and are associated with intestinal inflammatory responses^60–63^, providing a plausible link to the striking male-specific protection from DSS-induced colitis. The absence of similar lineage shifts in females may underlie their failure to gain colitis resistance despite harboring ever larger epithelial lipid stores. Sex hormones, which modulate tachykinin and receptor expression^64,65^, may be one contributing factor to this divergence and warrant further study.

Gut-innervating nociceptors regulate mucosal protection against dietary, microbial, and inflammatory insults^66–68^. These nociceptors control mucin secretion from goblet cells, to create a physical barrier between the epithelium and the luminal contents^69^. Recent evidence suggests that nociceptor-derived Substance P plays a crucial role in regulating mucosal protection during colitis and dysbiosis^66^. *Tac1* excision (which encodes both Substance P and NKA, the endogenous ligand for NK2R) from these neurons aggravates colitis^66^, whereas global NK2R ablation confers male-biased protection, suggesting that proinflammatory signaling through NK2R on immune or stromal cells may outweigh epithelial loss of tachykinin responsiveness during injury. In future efforts, conditional *Tacr2* deletion in distinct tissues as well as intestinal epithelial versus hematopoietic compartments will be essential to define the cell- and tissue-specific roles of NK2R.

Consistent with previous studies^57^, the HFD increased the fecal *Firmicutes*:*Bacteroidetes* (F:B) ratio after only one week, and this shift was reproduced in *Tacr2-/-* mice. We note here that our diets are matched for nutritional composition, differing only in fat and sugar sources and percent of total %kcal, limiting confounding effects of dietary fiber in our study. 18 operational OTUs were jointly influenced by genotype and diet; 12 belonged to the *Clostridiales* order [Families: *Lachnospiraceae* (5), *Ruminococcaceae* (3), and unclassified lineages (4)]. Members of *Lachnospiraceae* have been linked to constipation, whereas *Ruminococcaceae* abundance correlates with irritable bowel syndrome and type 2 diabetes^70^. Notably, the HFD expanded *Ruminococcus gnavus* (OTU575) and depleted *Akkermansia muciniphila* (OTU 1438), a combination associated with impaired barrier function through altered immune tolerance, short-chain fatty acid production and mucin layer maintenance^71–73^. Conversely, *Bifidobacterium pseudolongum* (OUT 9), which is often negatively associated with metabolic and inflammatory disorders^74–76^, was increased. Although this composite profile resembles that observed in inflammatory bowel disease, *Tacr2* deletion conferred resistance to DSS-induced colitis in males. These findings underscore the complexity of host-microbiota interactions and suggest that NK2R signaling modulates intestinal inflammation through mechanisms that extend beyond simple taxonomic shifts, potentially involving neuropeptide-dependent regulation of epithelial and immune function.

Collectively, our findings position NK2R at the intersection of metabolic and inflammatory pathways in the intestine, supporting the tachykinin-NK2R axis as highly tractable to therapeutic intervention. In principle, NK2R agonists could be leveraged to treat metabolic diseases including obesity, whereas NK2R antagonists may ameliorate mucosal inflammation in inflammatory bowel disease. The pronounced sexual dimorphism underscores the need to consider sex as a biological variable in the development of NK2R-targeted therapeutics. Based on our findings, we expect that delineating the precise mechanisms by which NK2R integrates neuronal, immune, and microbial cues to regulate lipid droplet dynamics and mucosal immunity in the intestine will yield important mechanistic insights and advance the development of targeted therapies for metabolic and inflammatory diseases.

## METHODS

### Animals

Male and female C57BL/6J mice (8-12 weeks old) were group-housed under a 12-h light/dark cycle with free access to food and water. Animals were fed either a low-fat control diet (CD; D14042701, Research Diets, New Brunswick, NJ; 73% kcal carbohydrate, 10% kcal fat, 17% kcal protein) or a Western-style high-fat diet (HFD; D12079B, Research Diets; 43% kcal carbohydrate, 40% kcal fat, 17% kcal protein). Body weights were measured weekly between 0900 and 1100 hours. All animal protocols were reviewed and approved by the Institutional Animal Care and Use Committee (IACUC) at The Scripps Research Institute.

### RNA extraction and RT-qPCR analysis

Mice were euthanized between 0900 and 1100 hours, and tissues were promptly collected and washed with ice-cold phosphate-buffered saline (PBS, 4°C) before snap-freezing in liquid nitrogen. The small intestine was washed with chilled PBS, opened longitudinally on ice, and the contents were removed. Intestinal epithelium was isolated by scraping with glass slides from the underlying submucosal layer and snap-frozen in liquid nitrogen. Samples were stored at −80°C until processing. Tissues were homogenized in TRIzol reagent, followed by phase separation with 1-bromo-3-chloropropane via centrifugation (8000g, 4°C). The upper aqueous phase was collected, and total RNA was purified using an RNeasy Mini Kit (Cat. No. 74104, Qiagen, Redwood City, CA) per the manufacturer’s instructions. cDNA synthesis was performed using iScript Reverse Transcription Supermix (Cat. No. 1708840, Bio-Rad, Hercules, CA). Quantitative PCR was conducted using SsoAdvanced Universal SYBR Green Supermix (Bio-Rad) according to the manufacturer’s protocols. Gene expression was normalized to reference housekeeping genes (*Actb*, *Rplp0*, or *Hprt*). RNA integrity was preserved by sanitizing all surfaces with 70% ethanol and RNase inhibitor (RNase OUT, G-Biosciences, St. Louis, MO). Data normalization was performed against either the small intestine or *Vil1* expression. Primer sequences are listed in the Supplementary Table 1.

### Immunohistochemistry

The proximal small intestine was excised and flushed with ice-cold modified Bouin’s fixative, then fixed in 10% buffered formalin for 24 hours at room temperature. Tissue was embedded and frozen in optimal cutting temperature (OCT) compound (Fisher Healthcare, Chino, CA) on dry ice. Sections (16 μm) were cut using a cryostat (Leica), mounted onto charged glass slides, permeabilized with 0.025% Triton X-100 in TBS, and blocked with 10% normal donkey serum and 1% bovine serum albumin (BSA) in TBST (Millipore Sigma, Cat. No. 566460). Sections were incubated with primary rabbit antibody against Neurokinin 2 Receptor (NK2R; 20 µg/mL; Cat. No. ATR-002, Alomone Labs, Jerusalem, Israel), followed by incubation with AlexaFluor 488-conjugated donkey anti-rabbit secondary antibody (Cat. No. A-21206, Thermo Fisher Scientific). Sections were washed, mounted with ProLong Gold Antifade reagent containing DAPI (Thermo Fisher Scientific) for nuclear staining, and imaged using a Nikon A1 confocal microscope with a 60x objective at room temperature. Image processing was conducted using ImageJ software version 2.0.0 (NIH, Bethesda, MD).

### CRISPR-Cas9 gene editing

Two single-guide RNAs (sgRNAs) were designed to target intronic regions flanking exon 2 of the *Tacr2* gene. A single-stranded DNA (ssDNA) repair template (721 nucleotides) containing homology arms (80 nucleotides each) flanking exon 2 and a LoxP sequence within the intronic regions was synthesized. Following Cas9-mediated DNA cleavage and homologous recombination using the ssDNA repair template, loxP sites were integrated flanking exon 2 to generate *Tacr2*^fl/fl^ mice. Non-homologous end joining after Cas9 cleavage resulted in the deletion of exon 2, generating *Tacr2*^−/−^ mice. Constructs (1 ng/µL) were microinjected into fertilized eggs to produce genetically modified mice, which were validated by sequencing. Genotyping primer sequences are listed in the Supplementary Table 2.

### Glucose Tolerance Test

Mice were fasted for 12 h prior to glucose tolerance testing. Fasted mice received an intraperitoneal injection of glucose solution (10% w/v) at a dose of 1 g glucose per kg body weight. Blood glucose concentrations were measured from tail blood samples at 0, 15, 30, 45, 60, and 120 min post-injection using a commercial handheld glucometer.

### RNA sequencing and analysis

RNA from intestinal epithelium was extracted as described above. Total RNA samples were prepared into RNA-Seq libraries using the NEBNext® Ultra II Directional RNA Library Prep Kit for Illumina following the manufacturer’s recommended protocol. Briefly, for each sample, 200 ng total RNA was polyA selected, and converted to double-stranded cDNA followed by fragmentation and ligation of sequencing adapters. The libraries were then PCR amplified 12 cycles using barcoded PCR primers, purified, and size selected using AMPure XP Beads before loading onto an Illumina NextSeq 2000 for 100 base single-read sequencing. Raw sequencing reads were quality-assessed using FastQC, and adapters were removed using Cutadapt. Trimmed reads were aligned to the mouse reference genome (ENSEMBL GRCm38) using STAR aligner version 2.6.1d^77^. The distribution of mapped reads across genomic features was analyzed with RSeQC v3.0.1. Differential gene expression analysis was performed using DESeq2 (v2.11.40.8) implemented in R (v4.2.2) via the Galaxy platform. Genes were considered significantly differentially expressed at thresholds of adjusted P < 0.05 and absolute log_2_ fold-change >1.0. Gene ontology, pathway enrichment, and network analyses were conducted using ShinyGO v0.80^78^. Cell-type-specific deconvolution was performed using BisqueRNA v1.0.5 against a published single-cell RNA sequencing dataset profiling mouse small intestinal epithelium^79,80^. Raw and processed sequencing data will be deposited in the Gene Expression Omnibus (GEO).

### Radioactive Tracer Oil challenges

Mice were fasted overnight prior to testing. Fasted mice were pretreated with Poloxamer-407. 10 g of Poloxamer-407 were resuspended in 100 mL of 0.9% NaCl saline and stirred overnight at 4°C. 10 µL/g of body weight was administered by intraperitoneal injection right before the oil gavage with 3 μCi [3H] triolein. Blood was collected at time 0, 1, 2, and 4 h, and the plasma was separated by centrifugation. Radioactivity was measured by scintillation.

### C_16_-BODIPY absorption assay

Mice were fasted overnight before receiving an oral gavage of olive oil containing 2 µg/g body weight of fluorescently labeled fatty acid analog C_16_-BODIPY (4,4-difluoro-5,7-dimethyl-4-bora-3a,4a-diaza-s-indacene-3-hexadecanoic acid; Thermo Fisher, cat. no. D3821) administered at a volume of 10 µL/g body weight. Mice were euthanized 4 h post-gavage, and intestinal tissues were collected, snap-frozen, sectioned, and imaged as previously described^81^.

### Gastrointestinal transit assay

Gastrointestinal transit time was assessed by oral gavage of a semi-liquid Evans Blue dye suspension (5% Evans Blue, 0.5% methylcellulose in phosphate-buffered saline; 100 µL per mouse). Following gavage, mice were monitored individually, and the time to appearance of the first blue-colored fecal pellet was recorded. Measurements were conducted in mice maintained on a control diet or after 12 weeks on a high-fat diet (HFD), as previously described^82^.

### FPLC assay

Mice were fasted overnight prior to testing and pretreated with either lipoprotein lipase (LPL) inhibitor Tyloxapol (500 µg/g body weight), vehicle (PBS), the NK2R agonist GR64349 (10 mg/kg), or the NK2R antagonist GR159897 (10 mg/kg). Following pretreatment, mice received an oral gavage of olive oil. Blood plasma was collected 2 h post-gavage, separated by fast protein liquid chromatography (FPLC), and triglyceride levels were quantified across collected fractions.

### Fecal lipid extraction

Mice were housed in wire-bottom cages for 12 h to collect fecal pellets. Pellets were dried, weighed, and pulverized using a mortar and pestle, then incubated in a 2:1 (v/v) chloroform:methanol solution. Samples were vortexed for 1 min and centrifuged at 1,000g for 10 min. The lower organic phase was collected, dried, and the residual lipids were weighed. Lipid content was normalized to the initial dried fecal weight^83^.

### Feeding behavior (CLAMS)

Indirect calorimetry was performed using a computer-controlled, open-circuit system (Oxymax System) integrated into the Comprehensive Lab Animal Monitoring System (CLAMS; Columbus Instruments, Columbus, OH)^84,85^. Single-housed, acclimated mice were placed into clear respiratory chambers (20 × 10 × 12.5 cm) equipped with a water sipper tube, a food tray connected to a balance for continuous food intake measurement, and 16 photobeams arranged in two axes at 0.5-inch intervals to monitor motor activity. Room air was circulated through each chamber at a flow rate of 0.5 L/min, and exhaust air was sampled every 15 min for 1 min. Oxygen consumption (VO_2_) and carbon dioxide production (VCO₂) were measured using O_2_ and CO_2_ sensors (Columbus Instruments), and respiratory exchange ratio (RER) was calculated as VCO_2_/VO_2_. All metabolic parameters were normalized to lean body mass, which was determined by EchoMRI analysis.

### Dextran sulfate sodium (DSS) colitis model

Colitis was induced by providing mice with *ad libitum* access to drinking water containing dextran sulfate sodium (DSS; 3.0% w/v for males, 5.0% w/v for females; MP Biomedicals, cat. no. MFCD00081551) for 7 consecutive days. Mice were monitored daily for changes in body weight and clinical condition. Animals that lost more than 25% of their initial body weight were euthanized and excluded from further analysis. On day 7, all remaining mice were euthanized, and tissues were collected for downstream analyses^86^.

### Tissue scoring parameters

The intact colon was collected, flushed with chilled phosphate-buffered saline (PBS), and fixed overnight in 4% paraformaldehyde (PFA) in PBS. The distal 1 cm segment was cryosectioned, stained with hematoxylin and eosin (H&E), and imaged. Histological sections were scored based on epithelial integrity (0-4) and immune cell infiltration (0-4) for a maximum aggregate score of 0-8^87^. Epithelial integrity was scored as follows: 0, normal morphology; 1, loss of goblet cells; 2, loss of goblet cells in multiple regions; 3, loss of crypts; 4, loss of crypts in multiple regions. Immune cell infiltration was scored as follows: 0, no infiltration; 1, infiltration around the crypt base; 2, infiltration reaching the lamina muscularis; 3, extensive infiltration of the lamina muscularis with mucosal thickening and edema; 4, infiltration extending into the submucosa.

### Fecal protein extraction for Lipocalin-2 quantification

Fecal pellets were collected at the time of sacrifice and flash-frozen in liquid nitrogen. Approximately 100 mg of frozen feces per sample was weighed, suspended in 1 mL of 1% Tween-20 in PBS, and vortexed vigorously for 10 min. Samples were then centrifuged at 15,000g for 10 min at 4°C. The resulting supernatant was collected and diluted (1:50 for control samples, 1:2000 for DSS-treated samples) prior to measurement of Lipocalin-2 concentrations using a commercial mouse Lipocalin-2 ELISA kit (R&D Systems, cat. no. DY1857)^49^.

### Intestinal permeability assay

Mice were fasted beginning at 09:00 for 4 h prior to oral gavage with fluorescein isothiocyanate-dextran 4000 (FD4; Millipore Sigma, cat. no. 46944) at a dose of 0.6 mg/g body weight from a 100 mg/mL stock solution. Four hours after gavage, blood was collected via cardiac puncture into BD Microtainer tubes containing lithium heparin and maintained on ice. Plasma was isolated by centrifugation at 5,000g for 5 min at 4°C. Plasma from fasted mice that did not receive FD4 was used to generate a standard curve (0-10 µg/mL FD4), and plasma FD4 concentrations in experimental samples were quantified accordingly^88,89^.

### Microbiome 16S rRNA sequencing and microbial analysis

Fecal pellets were collected from mice, and genomic bacterial DNA was extracted using the DNeasy PowerLyzer PowerSoil Kit (Qiagen, cat. no. 12855) according to the manufacturer’s instructions. DNA concentration was quantified and samples were submitted to GENEWIZ (South Plainfield, NJ, USA) for 16S-EZ sequencing using Illumina paired-end 2 × 250 bp methodology. Raw sequence reads were processed with Cutadapt (v1.9.1) to remove adapter sequences and assigned to operational taxonomic units (OTUs) based on a 97% similarity threshold using VSEARCH (v1.9.1) and QIIME (v1.9.1). Differential microbial abundance analyses were performed using linear discriminant analysis effect size (LEfSe)^90^.

### Statistical analysis

Data are presented as mean ± SEM. The number of biological replicates is provided in the figure legends. Comparisons between two groups were performed using unpaired two-tailed Student’s t-tests. Repeated measures two-way ANOVA was used for comparisons across time points, and one-way or two-way ANOVA followed by Sidak’s or Tukey’s post hoc multiple comparisons tests, as appropriate, were used for analyses involving more than two groups. Differences in UniFrac distances were assessed using PERMANOVA. Statistical significance was defined as P < 0.05. Outliers were identified using Grubb’s test. All statistical analyses were performed using GraphPad Prism v10.3.

## Acknowledgements

This work was supported by seed funding from the Scripps Research Institute. P.T. was supported by National Institutes of Health (NIH) grants R01 DK142199 and R01 HL175773. P.P. was supported by the NIH T32 Immunology Training Program (5T32AI007244). C.L. was supported by a Dorris Scholar Award from the Dorris Neuroscience Center at The Scripps Research Institute. We thank members of Enrique Saez’s lab at Scripps Research for technical assistance, support and critical advice during the course of the studies. We also thank Dr. Xinglin Yang (Howard Hang Lab) for technical expertise and guidance on microbiome workflows. We are grateful to Ayub Khan and Aayushi Shah (Srinivasan Lab) for technical assistance with tissue collection, and to Dr. Amanda Roberts (Scripps Research Animal Models Core) for support with mouse experiments. We also acknowledge the resources and staff of the Scripps Research Genomics Core Facility for sequencing services. Elements of Figures 2a, 3b, 4b, 4k, 5f, and Extended Data Figure 6b were created with BioRender.com.

## FIGURES and FIGURE LEGENDS

**Extended Data Figure 1.**
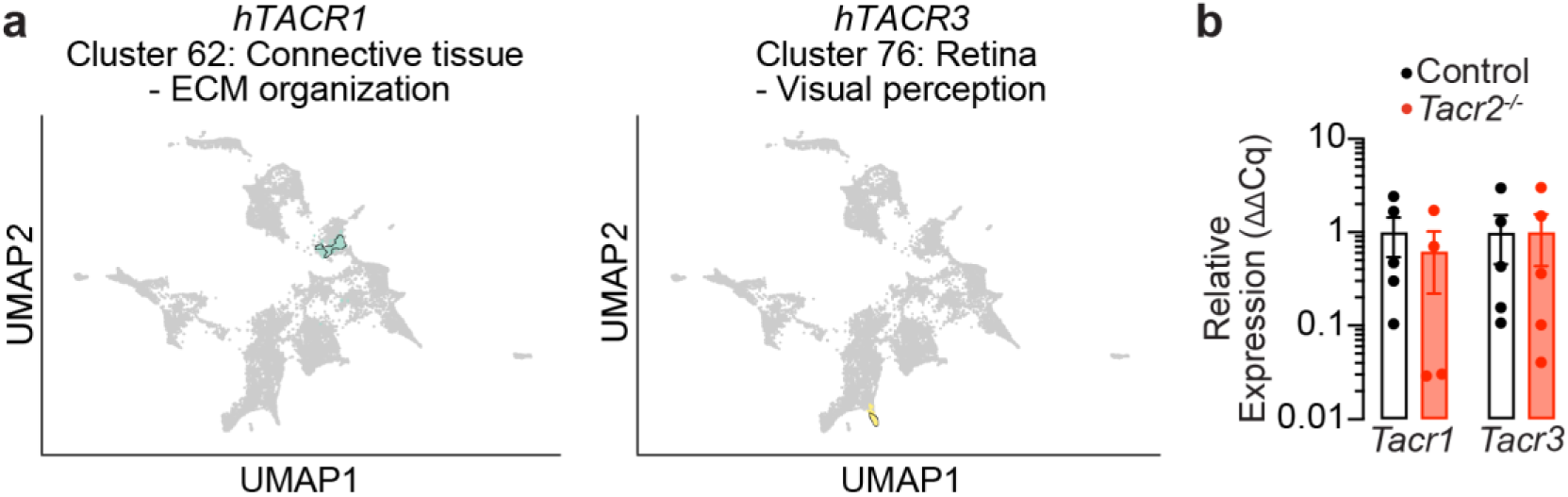
*Tacr1* and *Tacr3* expression in *Tacr2*^-/-^ male mice. **a,** UMAP visualization depicting gene clusters based on mRNA expression profiles from The Human Protein Atlas^44^ (HPA). The clusters containing *hTacr1* (cluster 62, comprising 389 genes) and *hTacr3* (cluster 76, comprising 175 genes) are highlighted. Functional annotation provided by HPA characterizes the shared specificity (Connective tissue or Retina) and biological function (ECM organization or Visual perception) of genes within the clusters. **b**, qPCR analysis of *Tacr1* and *Tacr3* mRNA expression in intestinal epithelial RNA extracted from male control and *Tacr2^-/-^* mice. n = 5 control and n = 4 *Tacr2^-/-^* mice for *Tacr1*, n = 5 per group for *Tacr3*.

**Extended Data Figure 2.**
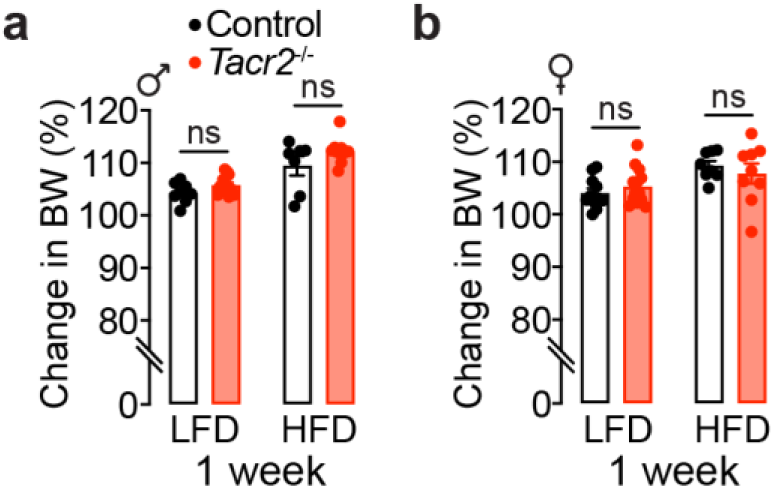
*Tacr2^-/-^*mice show no significant changes in body weight compared to wild-type controls under acute high-fat diet (HFD) challenge. **a,b**, Body weight changes from baseline (day 0) to day 7 in male (**a**) and female (**b**) control and *Tacr2^-/-^* mice fed a low-fat diet (LFD) or high-fat diet (HFD) for one week. n = 8 control (LFD), n = 12 *Tacr2^-/-^* (LFD), n = 7 control (HFD), and n = 9 *Tacr2^-/-^* (HFD) for (**a**); n = 11 control (LFD), n = 13 *Tacr2^-/-^*(LFD), n = 8 control (HFD), and n = 9 *Tacr2^-/-^* (HFD) for (**b**). Data were analyzed using two-way ANOVA with Holm-Sidak post hoc tests. Data are presented as mean ± SEM. ^ns^p > 0.05. Black bars indicate control mice; red bars indicate *Tacr2^-/-^* mice.

**Extended Data Figure 3.**
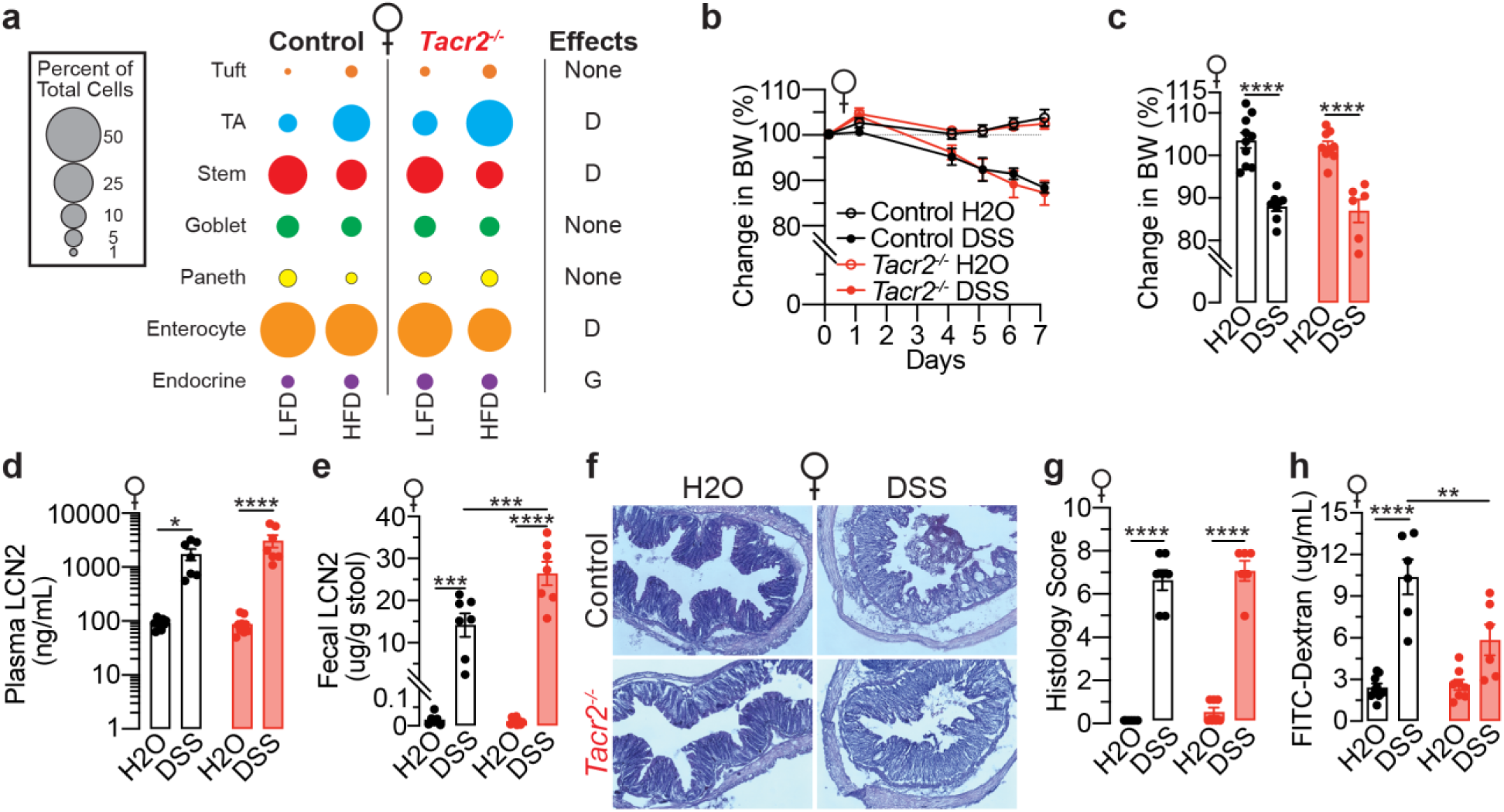
NK2R inhibition does not suppress DSS-induced weight loss and intestinal inflammation in female mice. **a**, Bulk RNA-Seq data were integrated with publicly available single-cell RNA-Seq datasets to estimate cell-type composition in the small intestine of female mice. Differences in estimated cell-type proportions were analyzed based on diet (D), genotype (G), or their interaction (I). **b**, Daily monitoring of body weights in female control and *Tacr2^-/-^* mice treated with DSS or vehicle over a 7-day period. n = 10 control (H_2_O), n = 9 control (DSS), n = 9 *Tacr2^-/-^*(H_2_O), and n = 7 *Tacr2^-/-^* (DSS). **c**, Comparison of body weight changes from baseline (day 0) to day 7, derived from the data shown in panel (**b**)**. d**,**e**, Lipocalin-2 (LCN2) concentrations in blood (**d**) and feces (**e**) from female control and *Tacr2^-/-^* mice after 7 days of DSS or vehicle treatment, as determined by ELISA. n = 9 control (H_2_O), n = 7 control (DSS), n = 9 *Tacr2^-/-^* (H_2_O), and n = 7 *Tacr2^-/-^* (DSS) for (**d**); n = 6 control (H_2_O), n = 7 control (DSS), n = 9 *Tacr2^-/-^*(H_2_O), and n = 7 *Tacr2^-/-^* (DSS) for (**e**). **f**, Colon tissues from female control and *Tacr2^-/-^* mice treated with DSS or vehicle for 7 days were collected, sectioned, and stained with hematoxylin and eosin (H&E). Representative images of stained colon sections are shown. **g**, Representative histological images (**f**) were evaluated and scored according to the degree of tissue damage. n = 6 control (H_2_O), n = 7 control (DSS), n = 7 *Tacr2^-/-^* (H_2_O), and n = 6 *Tacr2^-/-^*(DSS). **h**, Gut permeability in female control and *Tacr2^-/-^* mice following 7 days of DSS or vehicle treatment was assessed by oral gavage of 4 kDa FITC-dextran. Blood was collected 4 hours post-gavage, and FITC fluorescence was measured in plasma to quantify gut barrier integrity. n = 9 control (H_2_O), n = 6 control (DSS), n = 8 *Tacr2^-/-^*(H_2_O), and n = 6 *Tacr2^-/-^* (DSS).

**Extended Data Figure 4.**
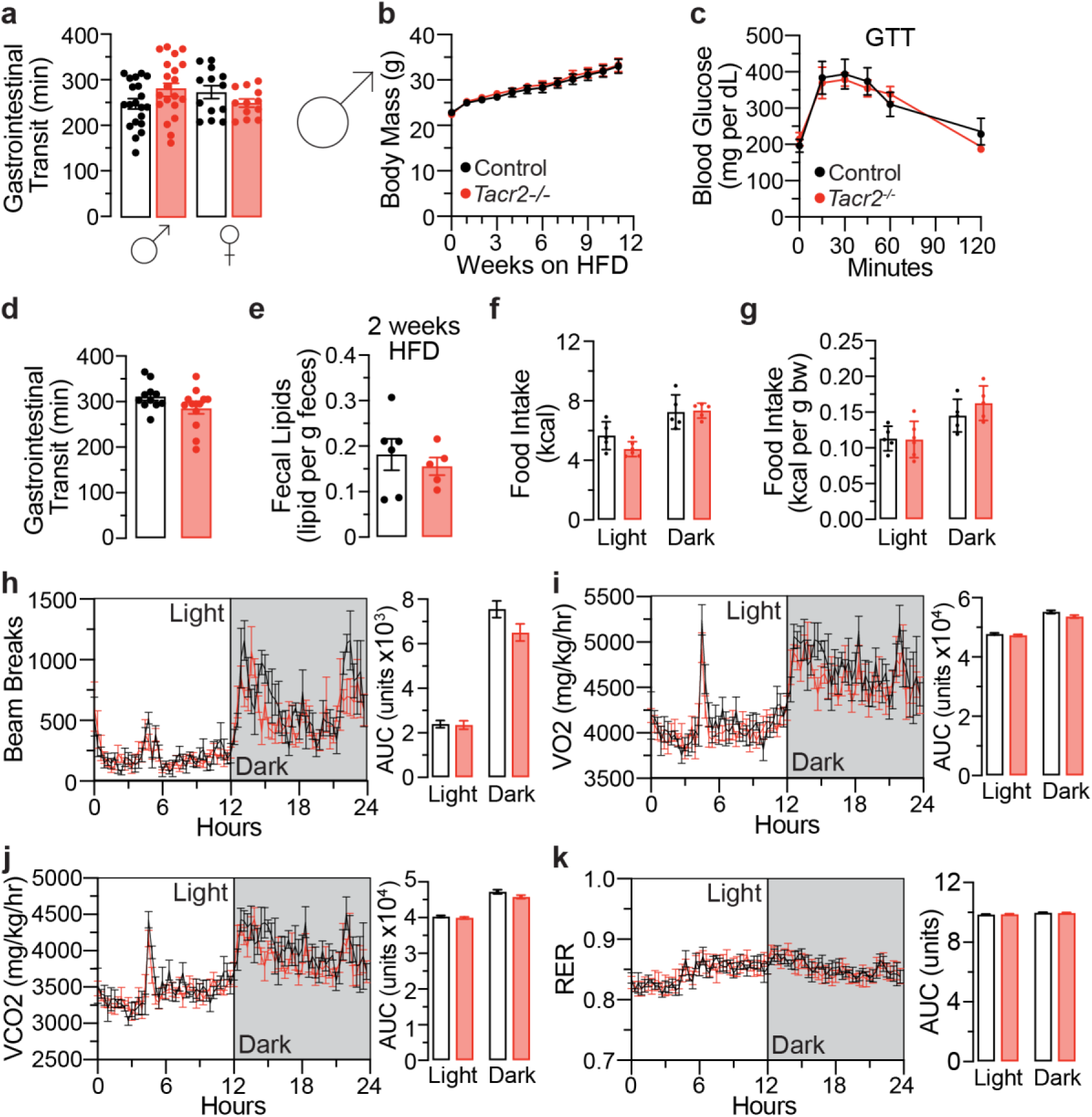
Male *Tacr2^-/-^* mice exhibit comparable gross phenotypes and feeding behaviors to control mice during chronic high-fat diet (HFD) feeding. a, Gastrointestinal transit time was measured in male and female control and *Tacr2^-/-^*mice on a low-fat diet (LFD). n = 11 control males, n = 13 *Tacr2^-/-^* males, n = 13 control females, and n = 12 *Tacr2^-/-^* females. **b**, The body weights of male control and *Tacr2^-/-^*mice (8-12 weeks old) were monitored weekly over a 10-week period while maintained on a HFD. n = 7 control and n = 9 *Tacr2^-/-^* mice. **c**, Glucose tolerance test (GTT) was performed with 10-week-old male control (n = 6) and *Tacr2^-/-^* (n = 5) mice. **d**, Gastrointestinal transit time was measured in male control (n = 11) and *Tacr2^-/-^* (n = 13) mice on a HFD. **e**, Fecal lipid content was quantified from stool samples collected at week 2 of HFD feeding in male control (n = 6) and *Tacr2^-/-^*(n = 5) mice. At week 10 of HFD treatment, male control and *Tacr2^-/-^*mice were placed in metabolic chambers, and food intake (**f**,**g**), locomotion (**h**), VO_2_ (**i**), VCO_2_ (**j**), and respiratory exchange ratio (RER, **k**) were analyzed. n = 5 control males (Light), n = 6 *Tacr2^-/-^* males (Light), n = 5 control males (Dark), n = 5 *Tacr2^-/-^* males (Dark) for (**f,g**); n = 4 control males (Light), n = 6 *Tacr2^-/-^* males (Light), n = 4 control males (Dark), n = 6 *Tacr2^-/-^* males (Dark) for (**h,i,j,k**). Data were analyzed using two-way ANOVA with repeated measures (**b,c**) or Holm-Sidak post hoc analysis (**f**-**k**), or by unpaired two-tailed Student’s t-test (**a**,**d**,**e**). Data are shown as mean ± SEM; ^ns^p>0.05.

**Extended Data Figure 5.**
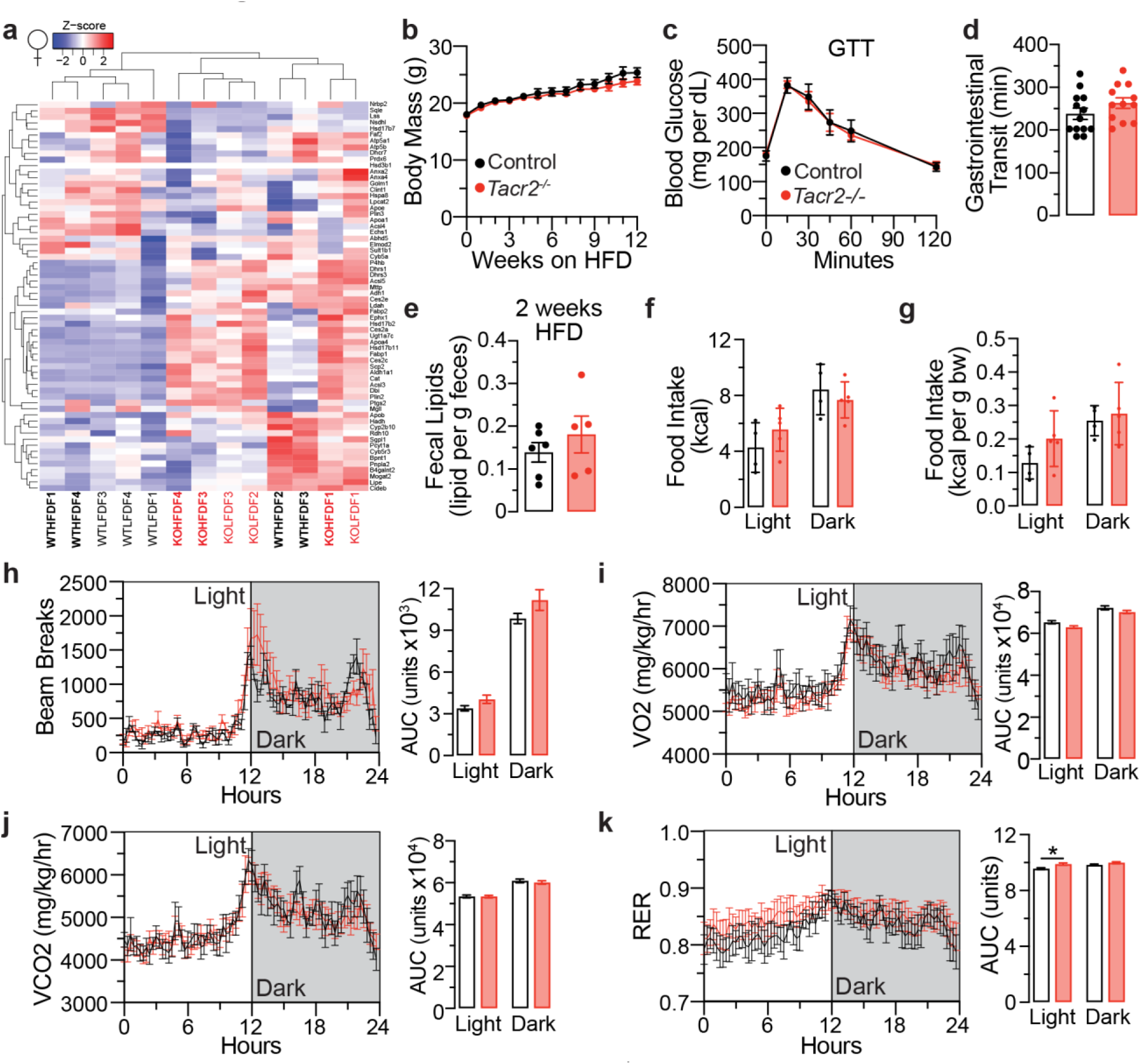
Female *Tacr2^-/-^* mice exhibit comparable gross phenotypes and feeding behaviors to control mice during chronic high-fat diet (HFD) feeding. **a**, Heat maps display Z-scores of normalized expression levels for genes encoding lipid droplet-associated proteins in female mice across experimental conditions. Hierarchical clustering of individual samples (top dendrogram) and genes (left dendrogram) is shown, with branch lengths reflecting the similarity between clusters. Wild-type control (WT) samples are labeled in black text, *Tacr2^-/-^* (KO) samples in red text, and samples from mice on a high-fat diet (HFD) are indicated in bold. **b**, The body weights of female control and *Tacr2^-/-^* mice (8-12 weeks old) were monitored weekly over a 10-week period while maintained on a HFD. n = 8 control and n = 9 *Tacr2^-/-^*mice. **c**, Glucose tolerance test (GTT) was performed with 10-week-old female control (n = 5) and *Tacr2^-/-^* (n = 7) mice. **d**, Gastrointestinal transit time was measured in female control (n = 13) and *Tacr2^-/-^* (n = 12) mice on a HFD. **e**, Fecal lipid content was quantified from stool samples collected at week 2 of HFD feeding in female control (n = 6) and *Tacr2^-/-^* (n = 5) mice. At week 10 of HFD treatment, female control and *Tacr2^-/-^*mice were placed in metabolic chambers, and food intake (**f**,**g**), locomotion (**h**), VO_2_ (**i**), VCO_2_ (**j**), and respiratory exchange ratio (RER, **k**) were analyzed. n = 4 control females (Light), n = 5 *Tacr2^-/-^* females (Light), n = 4 control females (Dark), n = 5 *Tacr2^-/-^* females (Dark) for (**f,g**); n = 4 control females (Light), n = 6 *Tacr2^-/-^* females (Light), n = 4 control females (Dark), n = 6 *Tacr2^-/-^* females (Dark) for (**h,i,j,k**). Data were analyzed using two-way ANOVA with repeated measures (**b**,**c**) or Holm-Sidak post hoc analysis (**f**-**k**), or by unpaired two-tailed Student’s t-test (**d**,**e**). Data are shown as mean ± SEM; *p<0.05, ^ns^p>0.05.

**Extended Data Figure 6.**
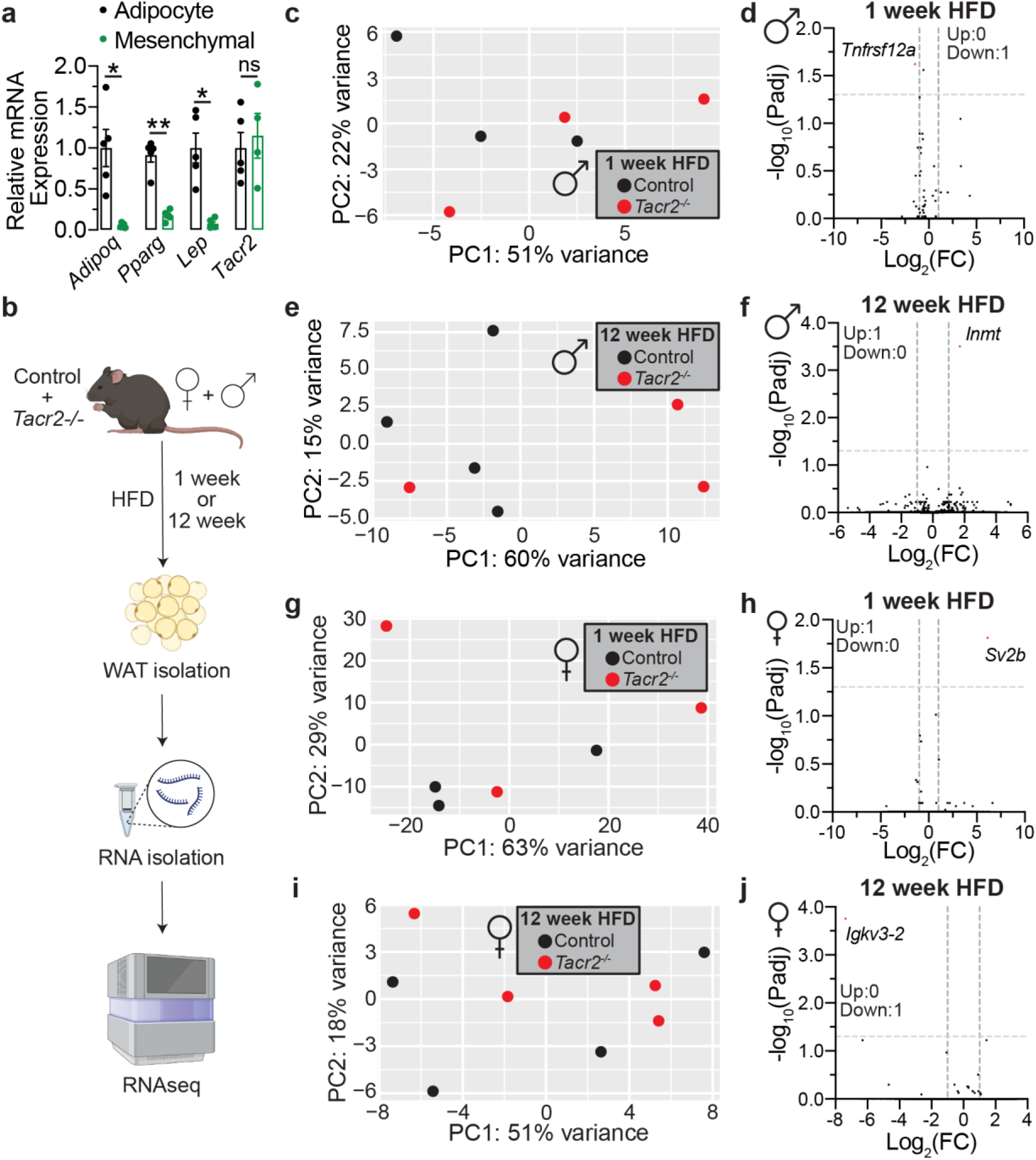
*Tacr2^-/-^*mice show similar gene expression profiles in perigonadal white adipose tissues. **a**, qPCR analysis of *Tacr2, Adipoq, Pparg, and Leptin* mRNA expression in adipocyte and mesenchymal cell fractions isolated from wild-type mice (n = 5 biological replicates). *Adipoq, Pparg, and Leptin* serve as adipocyte marker genes. **b**, Schematic of the experimental workflow. Control and *Tacr2^-/-^*mice were fed a high-fat diet (HFD) for either 1 or 12 weeks, followed by sacrifice and tissue collection. Perigonadal white adipose tissue (WAT) was harvested, total RNA was extracted, and samples were subjected to RNA sequencing. **c**,**e**,**g**,**i**, Principal component analysis (PCA) of RNA-Seq data from male (**c**,**e**) and female (**g**,**i**) control and *Tacr2^-/-^* mice following HFD feeding for 1 or 12 weeks. n = 3 control (1 week HFD), n = 4 control (12 week HFD), n = 3 *Tacr2^-/-^*(1 week HFD), n = 3 *Tacr2^-/-^* (12 week HFD) for (**c**,**e**); n = 3 control (1 week HFD), n = 4 control (12 week HFD), n = 3 *Tacr2^-/-^* (1 week HFD), n = 4 *Tacr2^-/-^*(12 week HFD) for (**g**,**i**). **d**,**f**,**h**,**j**, Volcano plots of differentially expressed genes in male (**d**,**f**) or female (**h**,**j**) *Tacr2^-/-^* versus control mice under 1 or 12 weeks HFD conditions. Data were analyzed using unpaired two-tailed Student’s t-test. Data are shown as mean ± SEM; **p<0.01, *p<0.05, ^ns^p>0.05; black circles represent control mice, and red circles represent *Tacr2^-/-^*mice.

---

**Supplementary Table 1:**
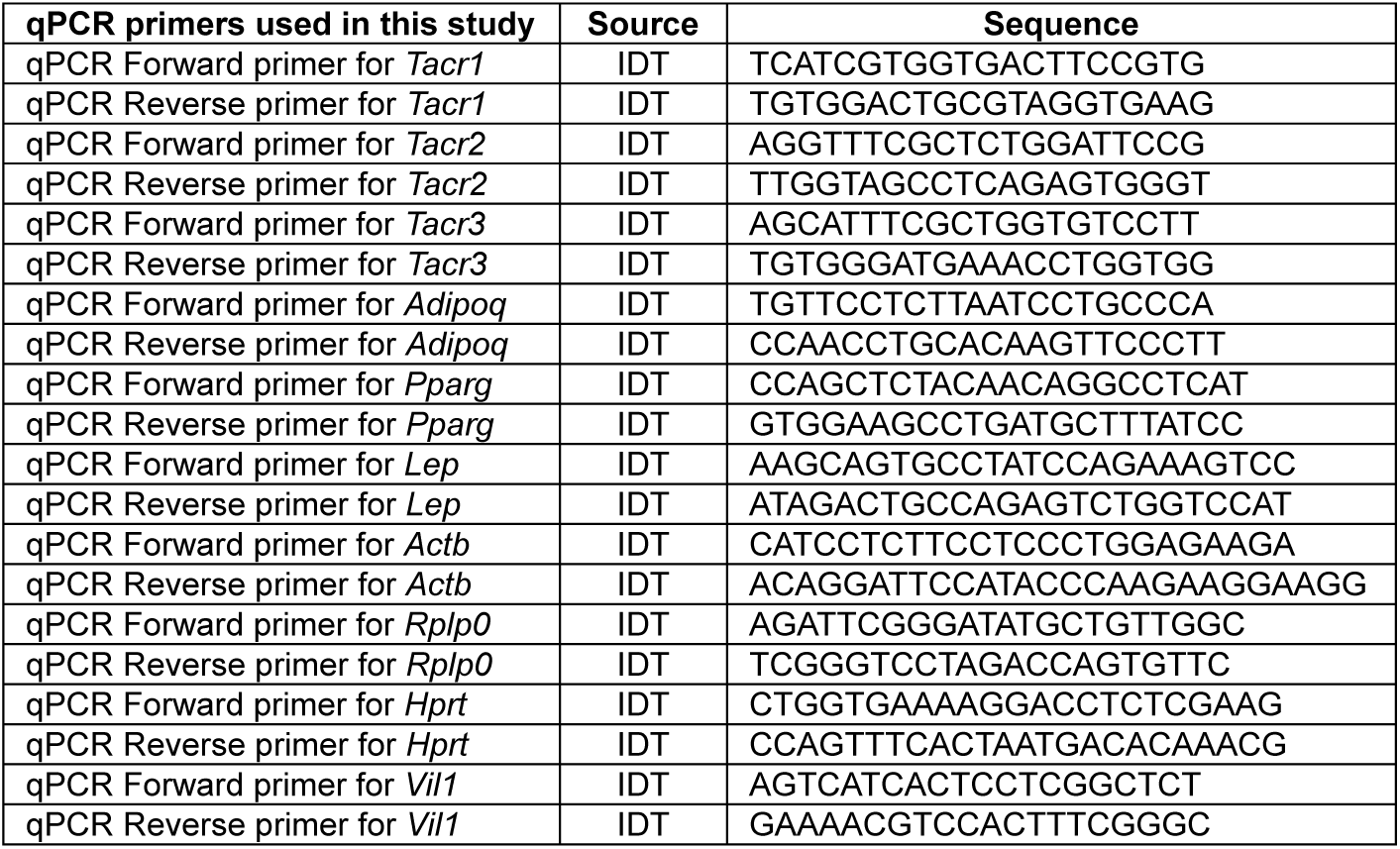
qPCR primers used in this study.

**Supplementary Table 2:**
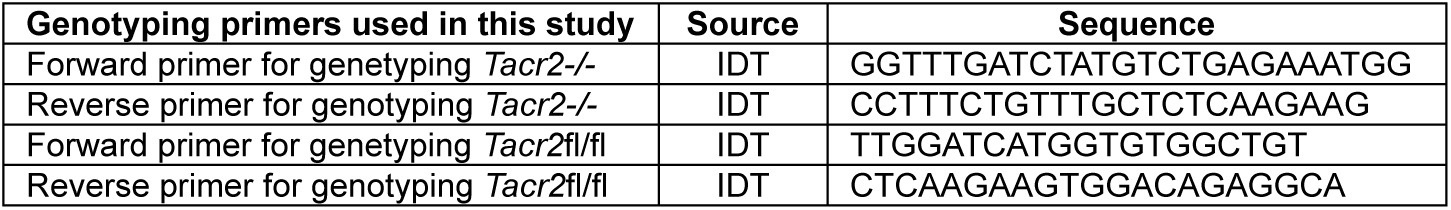
Genotyping primers used in this study.

## Notes

### Competing Interest Statement

The authors have declared no competing interest.

## References

1 Yoshinari, Y. et al. The sugar-responsive enteroendocrine neuropeptide F regulates lipid metabolism through glucagon-like and insulin-like hormones in Drosophila melanogaster. Nat Commun 12, 4818 (2021). 10.1038/s41467-021-25146-w

2 Palamiuc, L. et al. A tachykinin-like neuroendocrine signalling axis couples central serotonin action and nutrient sensing with peripheral lipid metabolism. Nat Commun 8, 14237 (2017). 10.1038/ncomms14237

3 Song, W., Veenstra, J. A. & Perrimon, N. Control of lipid metabolism by tachykinin in Drosophila. Cell Rep 9, 40–47 (2014). 10.1016/j.celrep.2014.08.060

4 Kuo, L. E. et al. Neuropeptide Y acts directly in the periphery on fat tissue and mediates stress-induced obesity and metabolic syndrome. Nat Med 13, 803–811 (2007). 10.1038/nm1611

5 Park, S. et al. Neuropeptide Y resists excess loss of fat by lipolysis in calorie-restricted mice: a trait potential for the life-extending effect of calorie restriction. Aging Cell 16, 339–348 (2017). 10.1111/acel.12558

6 Yin, Y. et al. AMPK-dependent modulation of hepatic lipid metabolism by nesfatin-1. Mol Cell Endocrinol 417, 20–26 (2015). 10.1016/j.mce.2015.09.006

7 Hou, L. et al. Neuropeptide ACP facilitates lipid oxidation and utilization during long-term flight in locusts. Elife 10 (2021). 10.7554/eLife.65279

8 Littlejohn, N. K., Seban, N., Liu, C. C. & Srinivasan, S. A feedback loop governs the relationship between lipid metabolism and longevity. Elife 9 (2020). 10.7554/eLife.58815

9 Nassel, D. R., Zandawala, M., Kawada, T. & Satake, H. Tachykinins: Neuropeptides That Are Ancient, Diverse, Widespread and Functionally Pleiotropic. Front Neurosci 13, 1262 (2019). 10.3389/fnins.2019.01262

10 Suvas, S. Role of Substance P Neuropeptide in Inflammation, Wound Healing, and Tissue Homeostasis. J Immunol 199, 1543–1552 (2017). 10.4049/jimmunol.1601751

11 Steinhoff, M. S., von Mentzer, B., Geppetti, P., Pothoulakis, C. & Bunnett, N. W. Tachykinins and their receptors: contributions to physiological control and the mechanisms of disease. Physiol Rev 94, 265–301 (2014). 10.1152/physrev.00031.2013

12 Zieglgansberger, W. Substance P and pain chronicity. Cell Tissue Res 375, 227–241 (2019). 10.1007/s00441-018-2922-y

13 Onaga, T. Tachykinin: recent developments and novel roles in health and disease. Biomol Concepts 5, 225–243 (2014). 10.1515/bmc-2014-0008

14 Hegron, A. et al. Therapeutic antagonism of the neurokinin 1 receptor in endosomes provides sustained pain relief. Proc Natl Acad Sci U S A 120, e2220979120 (2023). 10.1073/pnas.2220979120

15 Mantyh, P. W. Substance P and the inflammatory and immune response. Ann N Y Acad Sci 632, 263–271 (1991). 10.1111/j.1749-6632.1991.tb33114.x

16 Mishra, A. & Lal, G. Neurokinin receptors and their implications in various autoimmune diseases. Curr Res Immunol 2, 66–78 (2021). 10.1016/j.crimmu.2021.06.001

17 Martinez, A. N. & Philipp, M. T. Substance P and Antagonists of the Neurokinin-1 Receptor in Neuroinflammation Associated with Infectious and Neurodegenerative Diseases of the Central Nervous System. J Neurol Neuromedicine 1, 29–36 (2016). 10.29245/2572.942x/2016/2.1020

18 Li, Q. et al. Tachykinin NK1 receptor antagonist L-733,060 and substance P deletion exert neuroprotection through inhibiting oxidative stress and cell death after traumatic brain injury in mice. Int J Biochem Cell Biol 107, 154–165 (2019). 10.1016/j.biocel.2018.12.018

19 Vink, R. & van den Heuvel, C. Substance P antagonists as a therapeutic approach to improving outcome following traumatic brain injury. Neurotherapeutics 7, 74–80 (2010). 10.1016/j.nurt.2009.10.018

20 Donkin, J. J., Nimmo, A. J., Cernak, I., Blumbergs, P. C. & Vink, R. Substance P is associated with the development of brain edema and functional deficits after traumatic brain injury. J Cereb Blood Flow Metab 29, 1388–1398 (2009). 10.1038/jcbfm.2009.63

21 Gabrielian, L. et al. Substance P antagonists as a novel intervention for brain edema and raised intracranial pressure. Acta Neurochir Suppl 118, 201–204 (2013). 10.1007/978-3-7091-1434-6_37

22 Zhang, W. W., Wang, Y. & Chu, Y. X. Tacr3/NK3R: Beyond Their Roles in Reproduction. ACS Chem Neurosci 11, 2935–2943 (2020). 10.1021/acschemneuro.0c00421

23 Navarro, V. M. et al. The integrated hypothalamic tachykinin-kisspeptin system as a central coordinator for reproduction. Endocrinology 156, 627–637 (2015). 10.1210/en.2014-1651

24 Lederman, S. et al. Fezolinetant for treatment of moderate-to-severe vasomotor symptoms associated with menopause (SKYLIGHT 1): a phase 3 randomised controlled study. Lancet 401, 1091–1102 (2023). 10.1016/S0140-6736(23)00085-5

25 Depypere, H., Lademacher, C., Siddiqui, E. & Fraser, G. L. Fezolinetant in the treatment of vasomotor symptoms associated with menopause. Expert Opin Investig Drugs 30, 681–694 (2021). 10.1080/13543784.2021.1893305

26 Schank, J. R. Neurokinin receptors in drug and alcohol addiction. Brain Res 1734, 146729 (2020). 10.1016/j.brainres.2020.146729

27 Zhang, W. W. et al. Tachykinin receptor 3 in the lateral habenula alleviates pain and anxiety comorbidity in mice. Front Immunol 14, 1049739 (2023). 10.3389/fimmu.2023.1049739

28 Hether, S., Misono, K. & Lessard, A. The neurokinin-3 receptor (NK3R) antagonist SB222200 prevents the apomorphine-evoked surface but not nuclear NK3R redistribution in dopaminergic neurons of the rat ventral tegmental area. Neuroscience 247, 12–24 (2013). 10.1016/j.neuroscience.2013.05.006

29 Nakamura, A. et al. Bidirectional regulation of human colonic smooth muscle contractility by tachykinin NK(2) receptors. J Pharmacol Sci 117, 106–115 (2011). 10.1254/jphs.11118fp

30 Meuchel, L. W. et al. Neurokinin-neurotrophin interactions in airway smooth muscle. Am J Physiol Lung Cell Mol Physiol 301, L91–98 (2011). 10.1152/ajplung.00320.2010

31 Lecci, A., Capriati, A. & Maggi, C. A. Tachykinin NK2 receptor antagonists for the treatment of irritable bowel syndrome. Br J Pharmacol 141, 1249–1263 (2004). 10.1038/sj.bjp.0705751

32 Delvalle, N. M. et al. Communication Between Enteric Neurons, Glia, and Nociceptors Underlies the Effects of Tachykinins on Neuroinflammation. Cell Mol Gastroenterol Hepatol 6, 321–344 (2018). 10.1016/j.jcmgh.2018.05.009

33 Torres, E. et al. Congenital ablation of Tacr2 reveals overlapping and redundant roles of NK2R signaling in the control of reproductive axis. Am J Physiol Endocrinol Metab 320, E496–E511 (2021). 10.1152/ajpendo.00346.2020

34 Leon, S. et al. Characterization of the Role of NKA in the Control of Puberty Onset and Gonadotropin Release in the Female Mouse. Endocrinology 160, 2453–2463 (2019). 10.1210/en.2019-00195

35 Mapp, C. E. et al. The distribution of neurokinin-1 and neurokinin-2 receptors in human central airways. Am J Respir Crit Care Med 161, 207–215 (2000). 10.1164/ajrccm.161.1.9903137

36 Kitamura, H., Kobayashi, M., Wakita, D. & Nishimura, T. Neuropeptide signaling activates dendritic cell-mediated type 1 immune responses through neurokinin-2 receptor. J Immunol 188, 4200–4208 (2012). 10.4049/jimmunol.1102521

37 Ohtake, J. et al. Neuropeptide signaling through neurokinin-1 and neurokinin-2 receptors augments antigen presentation by human dendritic cells. J Allergy Clin Immunol 136, 1690–1694 (2015). 10.1016/j.jaci.2015.06.050

38 Vannucchi, M. G. & Faussone-Pellegrini, M. S. NK1, NK2 and NK3 tachykinin receptor localization and tachykinin distribution in the ileum of the mouse. Anat Embryol (Berl) 202, 247–255 (2000). 10.1007/s004290000106

39 Holzer, P. & Holzer-Petsche, U. Tachykinin receptors in the gut: physiological and pathological implications. Curr Opin Pharmacol 1, 583–590 (2001). 10.1016/s1471-4892(01)00100-x

40 Lecci, A., Capriati, A., Altamura, M. & Maggi, C. A. Tachykinins and tachykinin receptors in the gut, with special reference to NK2 receptors in human. Auton Neurosci 126-127, 232–249 (2006). 10.1016/j.autneu.2006.02.014

41 Renzi, D., Pellegrini, B., Tonelli, F., Surrenti, C. & Calabro, A. Substance P (neurokinin-1) and neurokinin A (neurokinin-2) receptor gene and protein expression in the healthy and inflamed human intestine. Am J Pathol 157, 1511–1522 (2000). 10.1016/S0002-9440(10)64789-X

42 Sass, F. et al. NK2R control of energy expenditure and feeding to treat metabolic diseases. Nature 635, 987–1000 (2024). 10.1038/s41586-024-08207-0

43 Karlsson, M. et al. A single-cell type transcriptomics map of human tissues. Sci Adv 7 (2021). 10.1126/sciadv.abh2169

44 Uhlen, M. et al. Proteomics. Tissue-based map of the human proteome. Science 347, 1260419 (2015). 10.1126/science.1260419

45 Jew, B. et al. Accurate estimation of cell composition in bulk expression through robust integration of single-cell information. Nat Commun 11, 1971 (2020). 10.1038/s41467-020-15816-6

46 Lee, J. C. et al. Obesogenic diet-induced gut barrier dysfunction and pathobiont expansion aggravate experimental colitis. PLoS One 12, e0187515 (2017). 10.1371/journal.pone.0187515

47 Liu, T. C. et al. Western diet induces Paneth cell defects through microbiome alterations and farnesoid X receptor and type I interferon activation. Cell Host Microbe 29, 988–1001 e1006 (2021). 10.1016/j.chom.2021.04.004

48 Chassaing, B., Aitken, J. D., Malleshappa, M. & Vijay-Kumar, M. Dextran sulfate sodium (DSS)-induced colitis in mice. Curr Protoc Immunol 104, 15 25 11–15 25 14 (2014). 10.1002/0471142735.im1525s104

49 Chassaing, B. et al. Fecal Lipocalin 2, a Sensitive and Broadly Dynamic Non-Invasive Biomarker for Intestinal Inflammation. PLOS ONE 7, e44328 (2012). 10.1371/journal.pone.0044328

50 Ma, X., Torbenson, M., Hamad, A. R., Soloski, M. J. & Li, Z. High-fat diet modulates non-CD1d-restricted natural killer T cells and regulatory T cells in mouse colon and exacerbates experimental colitis. Clin Exp Immunol 151, 130–138 (2008). 10.1111/j.1365-2249.2007.03530.x

51 van der Logt, E. M., Blokzijl, T., van der Meer, R., Faber, K. N. & Dijkstra, G. Westernized high-fat diet accelerates weight loss in dextran sulfate sodium-induced colitis in mice, which is further aggravated by supplementation of heme. J Nutr Biochem 24, 1159–1165 (2013). 10.1016/j.jnutbio.2012.09.001

52 Cheng, L. et al. High fat diet exacerbates dextran sulfate sodium induced colitis through disturbing mucosal dendritic cell homeostasis. Int Immunopharmacol 40, 1–10 (2016). 10.1016/j.intimp.2016.08.018

53 Teixeira, L. G. et al. The combination of high-fat diet-induced obesity and chronic ulcerative colitis reciprocally exacerbates adipose tissue and colon inflammation. Lipids Health Dis 10, 204 (2011). 10.1186/1476-511X-10-204

54 Beilstein, F., Carrière, V., Leturque, A. & Demignot, S. Characteristics and functions of lipid droplets and associated proteins in enterocytes. Experimental Cell Research 340, 172–179 (2016). 10.1016/j.yexcr.2015.09.018

55 Bickerton, A. S. et al. Preferential uptake of dietary Fatty acids in adipose tissue and muscle in the postprandial period. Diabetes 56, 168–176 (2007). 10.2337/db06-0822

56 Vaughn, A. C. et al. Energy-dense diet triggers changes in gut microbiota, reorganization of gut-brain vagal communication and increases body fat accumulation. Acta Neurobiol Exp (Wars) 77, 18–30 (2017). 10.21307/ane-2017-033

57 Crawford, M., Whisner, C., Al-Nakkash, L. & Sweazea, K. L. Six-Week High-Fat Diet Alters the Gut Microbiome and Promotes Cecal Inflammation, Endotoxin Production, and Simple Steatosis without Obesity in Male Rats. Lipids 54, 119–131 (2019). 10.1002/lipd.12131

58 Gustafsson, J. K. & Johansson, M. E. V. The role of goblet cells and mucus in intestinal homeostasis. Nat Rev Gastroenterol Hepatol 19, 785–803 (2022). 10.1038/s41575-022-00675-x

59 Feng, X., Fluchter, P., De Tenorio, J. C. & Schneider, C. Tuft cells in the intestine, immunity and beyond. Nat Rev Gastroenterol Hepatol 21, 852–868 (2024). 10.1038/s41575-024-00978-1

60 Steele, S. P., Melchor, S. J. & Petri, W. A., Jr. Tuft Cells: New Players in Colitis. Trends Mol Med 22, 921–924 (2016). 10.1016/j.molmed.2016.09.005

61 Nowarski, R. et al. Epithelial IL-18 Equilibrium Controls Barrier Function in Colitis. Cell 163, 1444–1456 (2015). 10.1016/j.cell.2015.10.072

62 Nystrom, E. E. L. et al. An intercrypt subpopulation of goblet cells is essential for colonic mucus barrier function. Science 372 (2021). 10.1126/science.abb1590

63 Yi, J. et al. Dclk1 in tuft cells promotes inflammation-driven epithelial restitution and mitigates chronic colitis. Cell Death Differ 26, 1656–1669 (2019). 10.1038/s41418-018-0237-x

64 Villablanca, A. C. & Hanley, M. R. 17beta-estradiol stimulates substance P receptor gene expression. Mol Cell Endocrinol 135, 109–117 (1997). 10.1016/s0303-7207(97)00193-7

65 Pinto, F. M., Pintado, C. O., Pennefather, J. N., Patak, E. & Candenas, L. Ovarian steroids regulate tachykinin and tachykinin receptor gene expression in the mouse uterus. Reprod Biol Endocrinol 7, 77 (2009). 10.1186/1477-7827-7-77

66 Zhang, W. et al. Gut-innervating nociceptors regulate the intestinal microbiota to promote tissue protection. Cell 185, 4170–4189 e4120 (2022). 10.1016/j.cell.2022.09.008

67 Lai, N. Y. et al. Gut-Innervating Nociceptor Neurons Regulate Peyer’s Patch Microfold Cells and SFB Levels to Mediate Salmonella Host Defense. Cell 180, 33–49 e22 (2020). 10.1016/j.cell.2019.11.014

68 Di Giovangiulio, M. et al. The Neuromodulation of the Intestinal Immune System and Its Relevance in Inflammatory Bowel Disease. Front Immunol 6, 590 (2015). 10.3389/fimmu.2015.00590

69 Yang, D. et al. Nociceptor neurons direct goblet cells via a CGRP-RAMP1 axis to drive mucus production and gut barrier protection. Cell 185, 4190–4205 e4125 (2022). 10.1016/j.cell.2022.09.024

70 Jackson, M. A. et al. Gut microbiota associations with common diseases and prescription medications in a population-based cohort. Nat Commun 9, 2655 (2018). 10.1038/s41467-018-05184-7

71 Everard, A. et al. Cross-talk between Akkermansia muciniphila and intestinal epithelium controls diet-induced obesity. Proc Natl Acad Sci U S A 110, 9066–9071 (2013). 10.1073/pnas.1219451110

72 Yan, J., Sheng, L. & Li, H. Akkermansia muciniphila: is it the Holy Grail for ameliorating metabolic diseases? Gut Microbes 13, 1984104 (2021). 10.1080/19490976.2021.1984104

73 Henke, M. T. et al. *Ruminococcus gnavus*, a member of the human gut microbiome associated with Crohn’s disease, produces an inflammatory polysaccharide. Proceedings of the National Academy of Sciences 116, 12672–12677 (2019). doi:10.1073/pnas.1904099116

74 Tatsuoka, M., Shimada, R., Ohsaka, F. & Sonoyama, K. Administration of Bifidobacterium pseudolongum suppresses the increase of colonic serotonin and alleviates symptoms in dextran sodium sulfate-induced colitis in mice. Biosci Microbiota Food Health 42, 186–194 (2023). 10.12938/bmfh.2022-073

75 Ma, B. et al. Strain-specific alterations in gut microbiome and host immune responses elicited by tolerogenic Bifidobacterium pseudolongum. Sci Rep 13, 1023 (2023). 10.1038/s41598-023-27706-0

76 Bo, T. B. et al. Bifidobacterium pseudolongum reduces triglycerides by modulating gut microbiota in mice fed high-fat food. J Steroid Biochem Mol Biol 198, 105602 (2020). 10.1016/j.jsbmb.2020.105602

77 Dobin, A., Davis, C.A., Schlesinger, F., Drenkow, J., Zaleski, C., Jha, S., Batut, P., Chaisson, M., Gingeras, T.R. STAR: ultrafast universal RNA-seq aligner Bioinformatics 29, 15–21 (2012). 10.1093/bioinformatics/bts635

78 Ge, S. X., Jung, D. & Yao, R. ShinyGO: a graphical gene-set enrichment tool for animals and plants. Bioinformatics 36, 2628–2629 (2019). 10.1093/bioinformatics/btz931

79 Haber, A. L. et al. A single-cell survey of the small intestinal epithelium. Nature 551, 333–339 (2017). 10.1038/nature24489

80 Jew, B. et al. Accurate estimation of cell composition in bulk expression through robust integration of single-cell information. Nature Communications 11, 1971 (2020). 10.1038/s41467-020-15816-6

81 Zhang, P. et al. Lipin 2/3 phosphatidic acid phosphatases maintain phospholipid homeostasis to regulate chylomicron synthesis. The Journal of Clinical Investigation 129, 281–295 (2019). 10.1172/JCI122595

82 Anitha, M. et al. Intestinal Dysbiosis Contributes to the Delayed Gastrointestinal Transit in High-Fat Diet Fed Mice. Cellular and Molecular Gastroenterology and Hepatology 2, 328–339 (2016). 10.1016/j.jcmgh.2015.12.008

83 Kraus, D., Yang, Q. & Kahn, B. B. Lipid Extraction from Mouse Feces. Bio-protocol 5, e1375 (2015). 10.21769/BioProtoc.1375

84 Chang, Y. et al. Ablation of NG2 Proteoglycan Leads to Deficits in Brown Fat Function and to Adult Onset Obesity. PLOS ONE 7, e30637 (2012). 10.1371/journal.pone.0030637

85 Yang, G. et al. Central role of ceramide biosynthesis in body weight regulation, energy metabolism, and the metabolic syndrome. American Journal of Physiology-Endocrinology and Metabolism 297, E211–E224 (2009). 10.1152/ajpendo.91014.2008

86 Chassaing, B., Aitken, J. D., Malleshappa, M. & Vijay-Kumar, M. Dextran sulfate sodium (DSS)-induced colitis in mice. Curr Protoc Immunol 104, 15.25.11–15.25.14 (2014). 10.1002/0471142735.im1525s104

87 Erben, U. et al. A guide to histomorphological evaluation of intestinal inflammation in mouse models. Int J Clin Exp Pathol 7, 4557–4576 (2014).

88 Pedicord, V. A. et al. Exploiting a host-commensal interaction to promote intestinal barrier function and enteric pathogen tolerance. Science Immunology 1, eaai7732–eaai7732 (2016). doi:10.1126/sciimmunol.aai7732

89 Woting, A. & Blaut, M. Small Intestinal Permeability and Gut-Transit Time Determined with Low and High Molecular Weight Fluorescein Isothiocyanate-Dextrans in C3H Mice. Nutrients 10 (2018). 10.3390/nu10060685

90 Segata, N. et al. Metagenomic biomarker discovery and explanation. Genome Biology 12, R60 (2011). 10.1186/gb-2011-12-6-r60

